# States of dynamic connectivity flow in temporal multiplex networks: a case study in human epilepsy and postictal aphasia

**DOI:** 10.1101/2024.05.10.593507

**Authors:** Nicola Pedreschi, Agnès Trebuchon, Alain Barrat, Demian Battaglia

## Abstract

We present a methodological framework for analysing multi-frequency dynamic functional connectivity (dFC) in electrophysiological recordings. The approach characterises not only the magnitude of network reconfiguration over time, but also whether these changes are spatially random or, instead, spatially organised in ways that drive a slower reconfiguration of modular structure. We define a generative null model of multi-scale connectivity fluctuations that differ in their degree of spatiotemporal organisation, and we describe dFC flows through the joint assessment of (i) instantaneous reconfiguration speed and (ii) the extent and quality of ongoing modular reorganisation. Different combinations of these features delineate distinct “flow styles”, ranging from more liquid to more frozen dynamics. As a case study, we apply this framework to SEEG recordings from epileptic patients. We identify transitions between dynamic “allegiance states”, whose flow styles closely mirror those of the null model. Seizure onset is associated with a pronounced slowing of dFC-speed, while a specific post-ictal regime combines low speed with highly frozen allegiance, and aligns most strongly with clinician-annotated aphasia. These pilot results suggest that temporal multiplex network analyses can reveal transient, frequency-specific network regimes linked to symptom expression and offers a generalisable tool for dissecting fast network dynamics in intracranial recordings.

**Author Summary:** Cognitive functions rely on the brain’s capacity to continually reorganize interactions among neuronal populations. Capturing this flexibility requires describing not only average functional network structure but also how it evolves over time. Using time- and frequency-resolved coherence from intracranial recordings in epilepsy patients, we model dynamic functional connectivity as a temporal multiplex network spanning multiple frequency bands. We introduce a framework that distinguishes different “styles” of network reconfiguration—faster or slower, and more structured or more random.

## 1 Introduction

Effective neural computation relies on spatiotemporally rich dynamics of functional networks that, via their continuous reconfiguration enable information to flow, be transformed and stored across distributed neuronal populations [1–3]. Optimal cognitive performance appears to depend on both the flexibility and the coordination of system dynamics, allowing the system to reshape its functional coalitions without collapsing into either rigidity or disorder [4–11]. Loss of this balance is a hallmark of cognitive pathology: numerous disorders are associated with reduced dynamical flexibility accompanied by a breakdown of coordinated organisation [12–15]. Temporal lobe epilepsy provides a particularly compelling example [16, 17]: beyond alterations in time-averaged local and global coupling between neuronal populations [18, 19], abnormalities in the dynamics of functional connectivity have been shown to predict language impairment more accurately than static network descriptors [20].

Yet, despite accumulating evidence that the temporal evolution of functional interactions is central to cognition, methods for quantifying the flow of network reconfiguration remain comparatively limited. The field has invested heavily in describing *where* connections exist (the spatial topology of instantaneous or time-averaged frames), but has developed far fewer tools for capturing *how* and *when* configurations change over time. New approaches are therefore needed to characterise the “styles” of dynamic reorganisation that underlie flexible cognition and their subtle perturbation in disease.

Here, we develop a framework to characterise the flow of functional connectivity reconfiguration, capturing both its temporal variability and the degree of spatial organisation. We first validate the framework using illustrative toy models of temporal networks, and then apply it as a proof of concept to intracranial recordings from drug-resistant epileptic patients suffering from post-ictal aphasia. In this application, the spectrally rich nature of electrophysiological signals enables us to assess parallel reconfiguration flows across multiple frequency bands, giving us the opportunity to extend our framework to time-evolving multilayer (multiplex) networks [21–23]. Furthermore, variations in network dynamics can be directly linked to seizure evolution and to the transient language disruptions that occur throughout the extended recovery phase.

Previous work [9, 10] has characterised dynamic styles of network flow by quantifying the instantaneous rate of change of functional interactions, that is, the dynamic functional connectivity *speed* (dFC-speed; Figure 1.a). This metric, initially developed for fMRI, can be applied to any temporal network to track frame-to-frame variability. The superior temporal resolution of electrophysiological recordings, however, allows us to examine reconfiguration on a slower time scale than that of single-frame transitions. We can group sequences of frames into longer time windows (hereafter “slow windows”, to distinguish them from the “fast” windows used to estimate individual-frame functional connectivity). Within each slow window, we compute both the average dFC-speed and the tendency of nodes to remain in matching communities across rapid structural fluctuations. This latter property is captured by a *module allegiance* matrix defined for each slow window (Figure 1.b).

**Figure 1.**
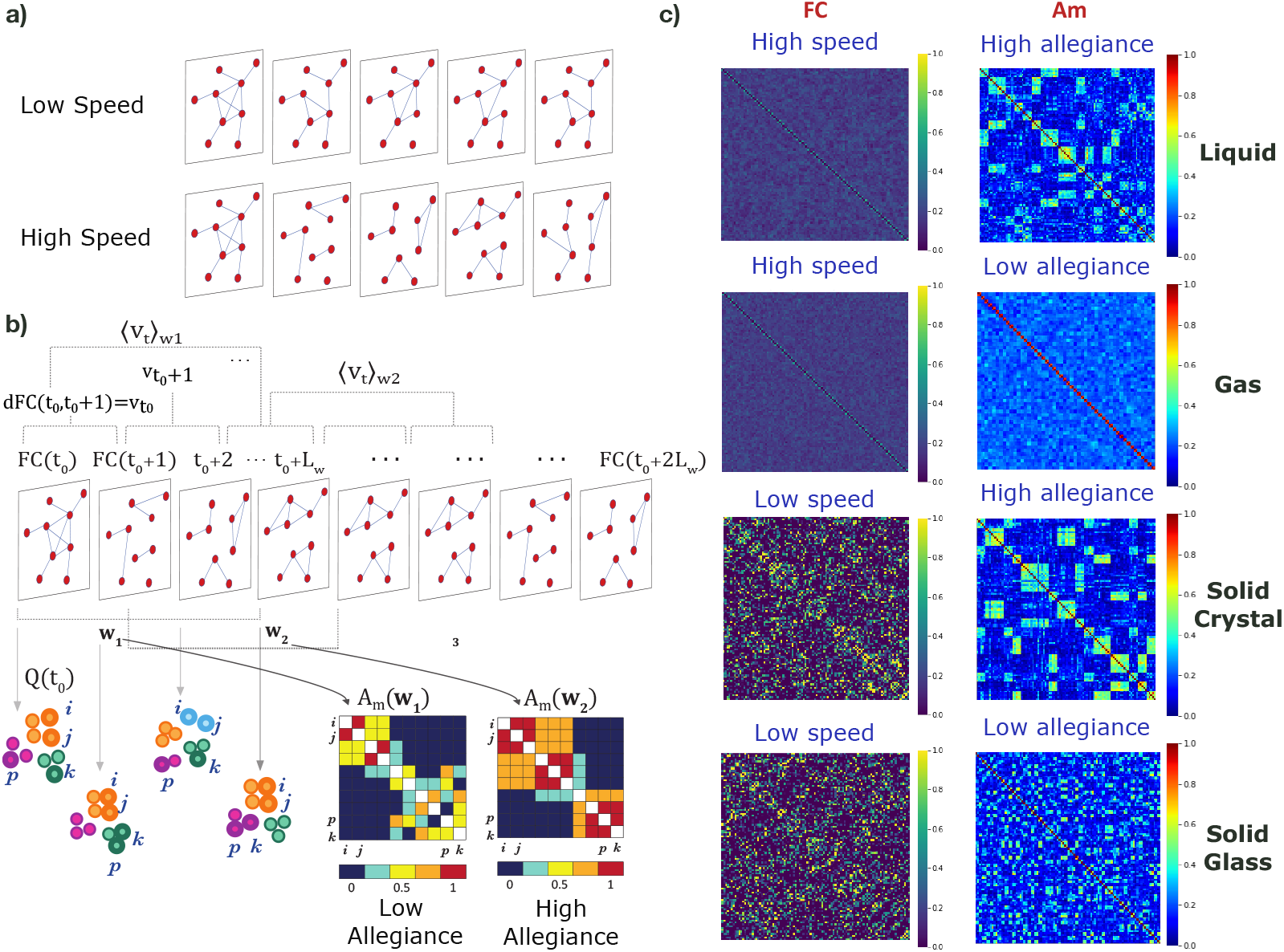
**a)** Cartoons of the snapshot-wise perspective of a slowly-varying temporal network (top) and of a rapidly-varying temporal network. **b)** From each pair of consecutive instantaneous FCs, we compute the rate of change, cosine distance, *dFC*(*t, t* + 1) (*v*_*t*_ for brevity) from one to the other, which we then average over a larger window *w*, including *L*_*w*_ time-frames. This procedure yields a window-resolved estimation of average *dFC-speed* ⟨*V*_*t*_⟩_*w*_ (schematic representation on the top). At each time point, we partition the network into its modules, and computing the associated modularity *Q*(*t*) (bottom-left cartoon). Within each window *w*, we compute the corresponding allegiance matrix *Am*(*w*), whose entries report the probability that each considered pair of network nodes (*i, j, p* and *k* in the cartoon) belong to the same network community in *w*. In **c)** we plot the average FC matrix (left) and average allegiance matrix 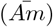 obtained for varying values of the *p*_*change*_ and *p*_*switch*_ parameters of our null model, corresponding to: high (*p*_*change*_ = 0.7) or low (*p*_*change*_ = 0.01) dFC-speed, and high (*p*_*switch*_ = 0.1) or low (*p*_*switch*_ = 0.7) allegiance.

To ground the interpretation of our metrics on intuitive examples, we first apply them to a generative null model of temporal networks based on a hierarchical extension of the Dynamic Stochastic Block Model (DSBM) [24, 25]. This model enables the simulation of temporal networks with tunable multi-level structure and dynamics, controlled by parameters that regulate the stability of individual links and of mesoscale modules. Within this model, we can generate distinct styles of network reconfiguration (Figure 1.c), which differ by their joint combination of speed and module allegiance.

We then apply the insights gained from this theoretical framework to empirical sEEG data. The null model provides a lens through which the complex physiological dynamics of the epileptic brain can be interpreted in terms of the idealized regimes identified earlier. For example, the pre-ictal baseline state maps onto a “liquid-like” regime (with simultaneously high allegiance and speed), whereas the long post-ictal recovery phase shows a progressive “liquefaction” of the “glassy-solid” dynamics (low speed and allegiance) observed during the seizure. We also find that transient “freezing” of dFC during recovery correlates with transient aphasia reported by clinicians, and that the flow of dFC across different frequency layers becomes more or less locked depending on the seizure stage.

Overall, our novel analytic framework goes beyond the simple localization of the epileptic focus. It offers a complementary perspective on how pathology alters the quality of network reconfiguration, potentially reflecting changes in low-level computational capabilities [13] that epilepsy may share with other conditions commonly observed in comorbidity [26].

## 2 Results

### 2.1 Null model of dynamic functional connectivity flows

We first aimed at generating surrogate dynamic network flows with controllable levels of temporal variability and spatial coordination in frame-to-frame fluctuations in order to benchmark our novel descriptive metrics. To do so we propose to use a hierarchical extension to the Dynamic Stochastic Block Model (DSBM) [24, 25, 27].

Initially, the network consists of *N* nodes distributed evenly among *n*_1_ first-level modules, which are then randomly assigned to *n*_2_ *< n*_1_ coarser, meso-scale second-level modules. Connections between nodes are governed by a classic stochastic block model: links between nodes in the same second-level module form with probability *p*_*in*_, while links between nodes in different second-level modules form with a lower probability, *p*_*out*_ *< p*_*in*_. This static network serves as the initial state of the temporal system.

Over time, two stochastic processes drive the network’s evolution. First, a global edge resampling mechanism randomly rewires the edges of the network at each time step, with each link being re-sampled with probability *p*_*change*_, or preserved with probability 1−*p*_*change*_. Second, a modular reorganization process stabilizes the hierarchy by occasionally reassigning entire first-level modules to new second-level communities. Specifically, a first-level module is reassigned to a different second-level module with probability *p*_*switch*_, while retaining its current assignment with probability 1 − *p*_*switch*_. When such a reassignment occurs, the edges of all nodes in the affected first-level module are adjusted to maintain the expected ratio of intra-to inter-module connection densities *p*_*in*_*/p*_*out*_ in the new configuration.

Such a hierarchical organisation allows two different aspects of the resulting flow of network reconfiguration to be tuned independently. The first is the reconfiguration *speed*, measured as the correlation distance between consecutive network frames (see Methods). This metric, introduced as dFC-speed in **(author?)** [9], quantifies the extent to which network structure changes from one frame to the next. Speed captures the overall number of altered links and increases when individual connections become less stable. In other words, regardless of the degree of coordination between link fluctuations, speed grows whenever one or more of the parameters *p*_*in*_, *p*_*out*_, *p*_*change*_ or *p*_*switch*_ are increased.

We then examine whether nodes switch between network modules independently, or in a manner coordinated with other nodes. This spatial coordination of temporal switching can be quantified using a *module allegiance* matrix, *Am*(*i, j*), which gives the probability that nodes *I* and *j* belong to the same module over time (Figure 1.b, bottom; see Methods for a detailed definition). The hierarchical structure of our toy model naturally favours the emergence of more or less persistent temporal communities—groups of nodes that remain densely and simultaneously interconnected throughout the network’s evolution. The extent of this persistence is controlled by the relative size of *p*_*switch*_, which governs coordinated module switching, with respect to *p*_*change*_, which sets the frequency of isolated link rewiring.

At each time frame, the network may exhibit a different modular partition. Fast reconfiguration can therefore affect modular structure in different ways. One possibility is that modules dissolve and reform almost randomly, with little memory of past organisation. In this case, the probability that any pair of nodes consistently belongs to the same transient module is low, and the resulting allegiance matrix contains only small, unstructured entries. Alternatively, reconfiguration may occur in a more organised fashion. Nodes can still move between modules, but do so in coordinated groups (the first-level modules), rather than independently. In this scenario, higher-valued entries emerge in the allegiance matrix, reflecting persistent co-assignment across time.

In this way, different settings of the null model parameters give rise to distinct *styles of network reconfiguration*, depending on the resulting levels of speed and module allegiance in the generated temporal network. Specifically (Figure 1.c, from top to bottom), we obtain:

- **liquid-like** flows, with high average speed and high allegiance
- **gas-like** flows, with high average speed but low allegiance
- **solid crystal-like** flows, with low speed and high allegiance
- **solid glass-like** flows, with low speed and low allegiance

These regimes are produced through ad hoc tuning of the model parameters, yet, as we show below, qualitatively similar regimes are observed in empirical dFC flows extracted from intracranial recordings. The naming of these flow styles provides an intuitive interpretative framework for understanding the combinations of speed and allegiance measured in real data, where reconfiguration is not controlled experimentally but reflects ongoing neural processes that shape dFC dynamics.

### 2.2 From intracranial SEEG recordings to a dynamic multiplex network

Having defined the different styles of temporal modular reconfiguration, we now turn to a real-world system in which such “gaseous”, “liquid” or “solid” dynamics may emerge along the dFC flow. In this work, we analyze with our approach intracranial recordings of 4 drugresistent epileptic patients (Figure 2a) with post-ictal aphasia, i.e., language production and comprehension impairments (Figure 2b). Detailed information on the recordings can be found in Table 1 and in the *Methods* section.

**Table 1:** Brain regions and relative electrodes count for the three patients.

| Brain area | Patient 1 | Patient 2 | Patient 3 |
| --- | --- | --- | --- |
| Frontal Operculum Cortex | 3 | 5 | 3 |
| Frontal Orbital Cortex | 7 | 5 | 3 |
| Frontal Pole | 6 | 34 | 23 |
| Inferior Frontal Gyrus, pars opercularis | 2 | 6 | 3 |
| Inferior Frontal Gyrus, pars triangularis | 1 | 2 | 0 |
| Inferior Temporal Gyrus, posterior division | 31 | 5 | 0 |
| Insular Cortex | 7 | 1 | 21 |
| Middle Frontal Gyrus | 5 | 5 | 25 |
| Middle Temporal Gyrus, anterior division | 5 | 17 | 5 |
| Middle Temporal Gyrus, posterior division | 16 | 16 | 0 |
| Parahippocampal Gyrus, anterior division | 8 | 5 | 5 |
| Parahippocampal Gyrus, posterior division | 3 | 2 | 0 |
| Planum Polare | 2 | 2 | 8 |
| Superior Temporal Gyrus, anterior division | 0 | 3 | 9 |
| Temporal Fusiform Cortex, anterior division | 5 | 1 | 0 |
| Temporal Fusiform Cortex, posterior division | 16 | 6 | 0 |
| Temporal Pole | 12 | 17 | 8 |
| unassigned | 7 | 20 | 36 |
| Cingulate Gyrus, posterior division | 4 | 0 | 4 |
| Inferior Temporal Gyrus, anterior division | 8 | 0 | 0 |
| Inferior Temporal Gyrus, temporooccipital part | 6 | 0 | 2 |
| Lingual Gyrus | 4 | 0 | 3 |
| Middle Temporal Gyrus, temporooccipital part | 6 | 0 | 5 |
| Precuneous Cortex | 2 | 0 | 2 |
| Superior Temporal Gyrus, posterior division | 2 | 0 | 5 |
| Temporal Occipital Fusiform Cortex | 7 | 0 | 1 |
| Angular Gyrus | 0 | 0 | 7 |
| Central Opercular Cortex | 0 | 0 | 5 |
| Cingulate Gyrus, anterior division | 0 | 0 | 6 |
| Heschl's Gyrus (includes H1 and H2) | 0 | 0 | 11 |
| Juxtapositional Lobule Cortex (formerly Supplem... | 0 | 0 | 3 |
| Lateral Occipital Cortex, superior division | 0 | 0 | 1 |
| Paracingulate Gyrus | 0 | 0 | 5 |
| Parietal Operculum Cortex | 0 | 0 | 3 |
| Planum Temporale | 0 | 0 | 8 |
| Postcentral Gyrus | 0 | 0 | 8 |
| Precentral Gyrus | 0 | 0 | 11 |
| Superior Frontal Gyrus | 0 | 0 | 4 |
| Superior Parietal Lobule | 0 | 0 | 1 |

**Figure 2.**
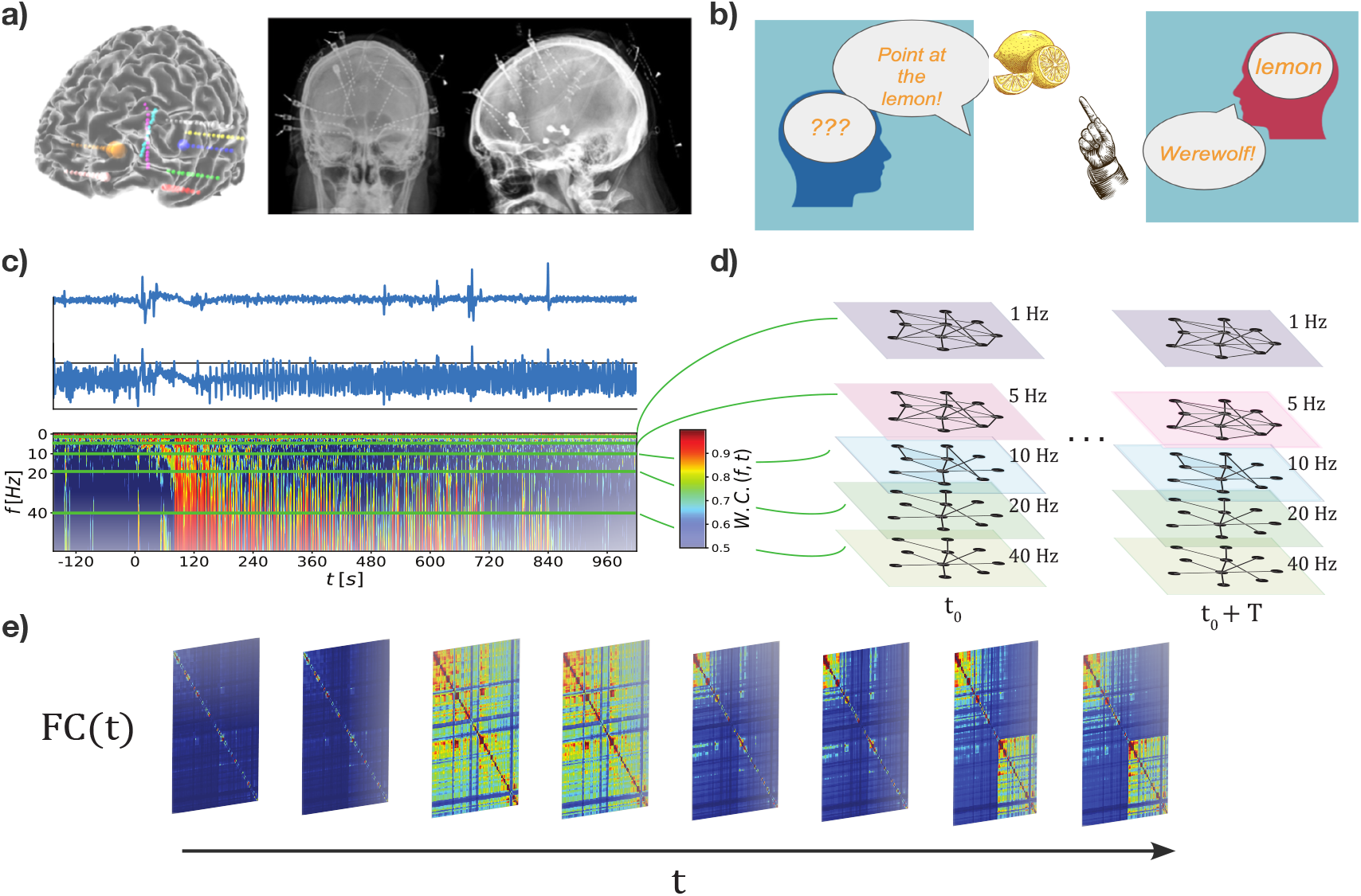
**a)** Representative 3D reconstruction and X-ray imaging of intracranial stereotactic EEG (sEEG) electrodes implanted in a patient suffering of pharmaco-resistant epilepsy for the sake of pre-surgical investigations and determination of the epileptogenic zone. Intracranial recordings are performed over long durations reaching several hours, including one or more seizure events, surrounded by pre-ictal and post-ictal periods. The positioning of multiple electrodes allows recording simultaneously activity from a variety of temporal, parietal and frontal brain regions (See Table S1) **b)** During the recovery phase following a seizure, transient impairments of the language function (aphasia) are reported by the clinicians. The cartoons represent examples of impairments that can be detected through the testing of, for instance, Wernicke’s aphasia (left) and anomia (right).**c)** Schematic representation of a recording as a dynamic multiplex network; from each pair of signals from different electrode contacts *A* and *B* (blue curves on the top left, examples for two representative channels) we compute the *Morlet wavelet coherence* at each time-stamp for a span of frequencies ranging from 1 to 80 Hz; we consider 5 frequencies of interest, 1, 5, 10, 20 and 40 Hz and follow their dynamics edges in the respective 5 layers (on the right) for each pair of regions. **d** We therefore end up with an instantaneous network for each time-stamp in each of the 5 layers. **e)** For each layer (frequency) we generate a series of instantaneous snapshots of the network, i.e., a stream of FC matrices (middle).

Different frequency bands are thought to participate in the processing of distinct aspects of language, and to contribute in a coordinated manner [28, 29]. Since neural interactions unfold simultaneously across multiple frequency bands, the system under study can naturally be represented not as a single evolving network but as a *multiplex* of frequency-specific layers. To capture these multilayer interactions, we compute the wavelet coherence [30] in five different frequency bands between each pair of recorded signals. To do so, we decompose signals in the time-frequency domain along the Morlet wavelet family (time–frequency power spectra are provided in Supplementary Figure S1). The wavelet coherence *W.C*.(*t, f*) (Figure 2.c, see Methods for proper definition) has values between 0 and 1. We compute the wavelet coherence *W.C*(*t, f*) for each pair of signals in 5 frequency bands peaked around 1Hz, 5Hz, 10Hz, 20Hz and 40Hz. The coherence values computed for each of the 5 reference frequencies represent, for each pair of signals, 5 time series of temporal edges between the same nodes *i* and *j* representing the sources of the signals. We therefore represent these data as a time-varying multi-layer weighted network, where the weight of a temporal edge (*i, j, t*)^*l*^ at time *t*, in layer *l* corresponds to the coherence in frequency band *l* of nodes *i* and *j* at time *t*. Since the nodes in all layers are the same, we refer to this object as a *temporal multiplex network*. Only significant links are included in the multiplex networks, where a significance threshold is determined in terms of coherence estimated with randomized signals. After signal down-sampling, we maintain a very precise temporal resolution, since, for the faster frequencies, we estimate a different, independent time-resolved network frame every ∼100ms (see *Materials and Methods*). A schematic representation of the extraction of the temporal multiplex from the raw data can be found in Figure 2d,e.

We first present a complete analysis for one representative seizure recording. Focusing on a single concrete example allows us to illustrate each step of the analysis pipeline in detail while keeping the exposition clear and concise. A systematic comparison across all available recordings—four seizures from three patients—is then provided in the final section. Although the sample size is necessarily small, as is common in SEEG studies, we note here that several qualitative patterns observed in the representative example are consistently replicated across the remaining recordings.

### 2.3 Layer-wise global network properties

We start our analysis by computing two global network measures on each layer as general indicators of connectivity fluctuations. Such measures are the instantaneous total network strength, namely the sum of edges’ weights at each time point, and the layer’s modularity, i.e., a quality function *Q*(*t*) ∈ [0, 1] which reflects the tendency of the network’s layer to be organised into different modules (communities; in the literature *Q* ≥ 0.2 for significant network partition into different modules/communities [31, 32]). See Methods for a detailed description of our the community detection approach.

In Figure 3.a, we show for each different frequency layer, the time series of the layer-wise instantaneous total strength *s*(*t*) (sum of all instant link weights) of the network (blue hues), and the instantaneous modularity *Q*(*t*) (orange hues). In Figure 3.a, for the 1Hz layer, the strength *s* (darkest blue line in the top plot) is persistently close to *N* = 117 (the size of the network), meaning that, in average, each node is connected to almost the entirety of the rest of the network with high values of coherence. This layer of the network is therefore particularly dense, associated to global slow oscillations of the entire system, which causes the values of *Q*(*t*) to be extremely low (darkest red line in the bottom plot). This suggests that the 1Hz layer has no modules to be retrieved at any given time.

**Figure 3.**
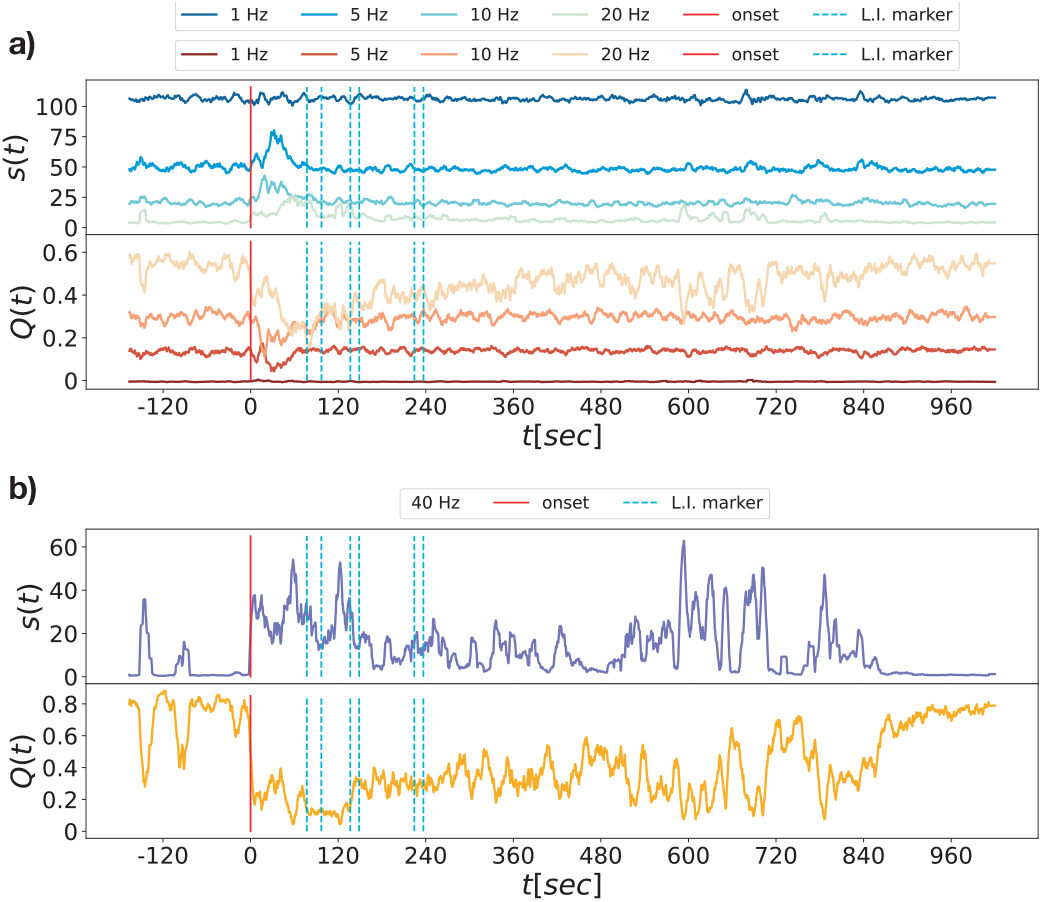
Strength and Modularity time series: a) We show, for a representative recording, time series of instantaneous total strength *s*(*t*) (top, blue colors) and modularity *Q*(*t*) (bottom, red colors) of four frequency bands (in progressively lighter colors from slower to larger frequencies, namely: the 1Hz, 5Hz, 10 Hz and 20Hz layers. In b), we show with greater detail, *s*(*t*) and *Q*(*t*) time-series for the highest-frequency layer at 40Hz. The vertical lines in the plots represent the clinical markers: the seizure onset in red, and the language dysfunction markers are represented by the cyan dotted lines. See Supporting Figure S2 for other recordings and Supporting Figure S3 for comparison between uni- and bipolar montages.

When increasing frequency (lines with progressively lighter colors in Figure 3.a), progressively richer dynamics emerge, partly due to lower connection densities. In the 5Hz layer, average strength is lower than at 1Hz, then exhibits a sharp rise at seizure onset followed by a rapid return to baseline, occurring even before any reported clinical marker. Modularity remains below *Q* ∼ 0.2 throughout the recording, yet shows a clear drop at seizure onset, coincident with the transient strength increase. Strength and modularity are anti-correlated across layers: for the 10Hz layer, both trends resemble those at 5Hz, with lower *s*(*t*) and higher *Q*(*t*), though the return to baseline occurs slightly later.

This frequency-dependent slowing of recovery is more evident at higher frequencies. In the 20Hz layer (Figure 3.a, lightest colors), both strength and modularity return to baseline markedly later, in fact after the last clinical marker—i.e., beyond the period during which clinicians report post-ictal symptoms. The 40Hz layer (Figure 3.b) shows the slowest recovery: here the seizure-induced perturbation persists longest, with baseline levels re-emerging only in the final minutes of the recording, well after seizure termination.

Analogous strength and modularity patterns across recordings are shown in Supplementary Figure **S2**. Qualitatively similar trends also appear when networks are constructed from bipolar signals (Supplementary Figure **S3**). We focus here on unipolar montages because their larger number of nodes, despite increased noise and uncorrected volume conduction, enhances the visibility of network structure in the higher-frequency layers (see *Discussion*).

### 2.4 Comparing network flows, beyond networks: dynamic Modular Allegiance analysis

As described above, we perform the layer-wise analysis at two time resolutions: a fast one, defined by individual network frames, and a slower one, defined by sliding windows of *L*_*w*_ = 200 time points (approximately 2.3 seconds). This double resolution allows us to characterize both the organization of single-frame networks (Figure 3) and the time-dependence of their frame-to-frame reconfiguration, since each window contains multiple frames and thus provides a localized “snippet” of evolving Functional Connectivity.

Within each slower window *w*, we compute the allegiance between all region pairs, quantifying the degree to which they shift across transient modules in a similar fashion. This yields a time-resolved stream of *allegiance matrices Am*(*w*), where *Am*_*ij*_(*w*) is the probability that regions *i* and *j* belong to the same module within that window (Figure 2.b, bottom). Remember that dynamic allegiance is evaluated on the slow time scale set by the window size *w*, unlike speed, which is evaluated on the fast frame-to-frame scale and only later averaged within each window. Sliding the window produces a temporally ordered sequence of allegiance matrices for each layer, capturing fast reconfiguration *within* windows and slower “dynamics of network dynamics” *across* windows. Following the intuition developed from the null model (Figure 1.c), the mean within-window allegiance expresses how random or structured the reconfiguration is within the considered slow window, while block structure reveals sets of regions flowing together in a more “liquid-like” manner, in contrast to the more “gas-like” random reassignment observed during low-allegiance periods.

The *style* of module reconfiguration may itself vary over time. To detect changes in this style, we compare allegiance matrices across windows via cosine similarity (see *Methods*). These similarities form the *dynamic Module Allegiance matrix* (*dA*), a time–time matrix analogous to the *dFC matrix* used in fMRI [9, 10] or to recurrence matrices tracking network-topology evolution [1, 2]. In our case, however, we compare not connectivity patterns but connectivity *flows*, summarized by the allegiance matrices; fast variations occur within windows, while slower changes across windows appear in the *dAm* structure. For each layer, the dynamic allegiance matrix *dAm*^*l*^ (Figure 4.a, 40Hz band) is computed as the cosine similarity between allegiance matrices from windows *t* and *t*^*∗*^. Each element *dAm*^*l*^(*t, t*^*∗*^) ranges from 0 (completely dissimilar) to 1 (identical). Block structure in *dAm*^*l*^ reveals epochs—or *allegiance states* [33]—during which allegiance matrices remain similar and the flow pattern is stable. We cluster the layer-wise tensor *Am*_*ij*_(*t*) via spectral clustering, using the *dAm* matrix as the precomputed similarity across time windows.

**Figure 4.**
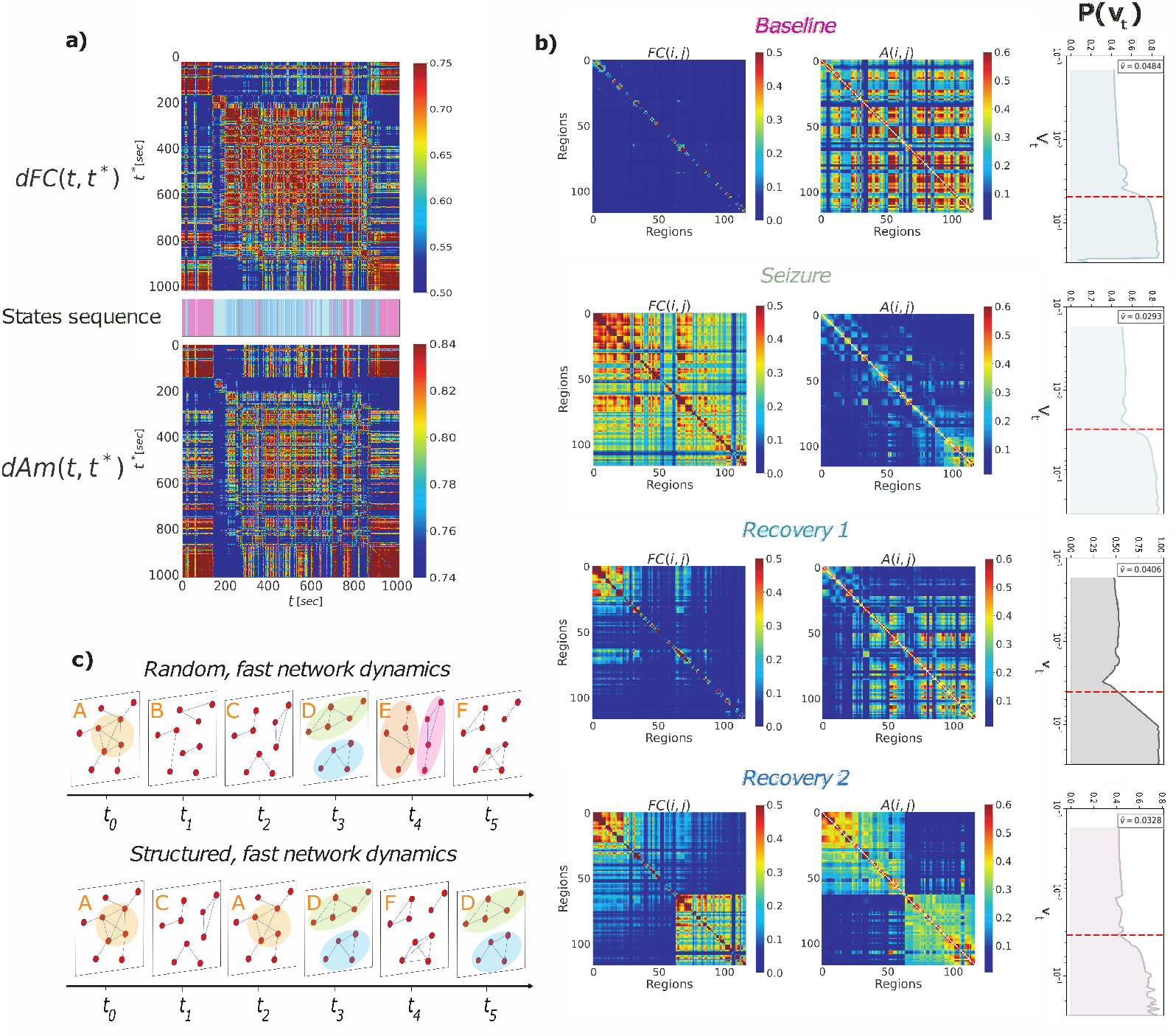
Allegiance dynamics and allegiance states: **a)** dynamic Functional Connectivity (dFC) matrix (top) and dynamic Allegiance matrix (dAm), bottom, computed for the 40Hz frequency band of the representative recording. **b)** Each row corresponds to one of four retrieved allegiance states, namely, Baseline, Seizure, Recovery 1 and Recovery 2, and the three plots on a row are, from left to right, the corresponding average Functional Connectivity matrix, the corresponding average modular allegiance matrix and the distribution of dFC-speed values measured within the corresponding state; the state-wise average value 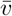 reported in Table 5 are annotated on the plots. In **c)** is a cartoon comparison of different flows of network dynamics that our pipeline allows us to capture: for instance we can distinguish a fast, random rewiring of the network and re-organization of the communities lying within it, from a more structured dynamics.

### 2.5 Modular allegiance states

In Figure 4.b we show the four allegiance states (clusters of time windows in *dAm*) identified for the 40Hz band of a representative patient. Analogous analyses for lower-frequency layers are shown in Supplementary Figures S4 and S5, while the same dFC and allegiance analyses for the 40 Hz layer in the other recordings are reported in Supplementary Figure S6. For each state, the left matrix displays the average Functional Connectivity (FC) computed over the windows assigned to that state (e.g., the *Baseline* state in the first row). This FC matrix shows generally low off-diagonal values, with small diagonal blocks of tightly connected neighboring contacts—consistent with the high modularity seen before seizure onset and at the end of the recording in Figure 3.b, and likely influenced by volume conduction in unipolar signals.

The right matrix shows the allegiance averaged over the same baseline windows 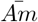. Its lattice-like block structure reveals multiple groups of nodes with high allegiance, including clear off-diagonal blocks not restricted to spatial neighbors. Thus, even during baseline, the allegiance structure is highly organized, indicating that within-window modular reconfiguration is not random. This raises the question of whether the underlying modular structure changes at all during baseline windows.

High allegiance values alone may simply indicate that the modular structure remains stable throughout a window. To distinguish a “ crystallized” partition—where nodes keep fixed module memberships—from a “liquid” reconfiguration—where nodes change modules in a coordinated but non-random manner—*dAm* is not sufficient, as previously discussed; we must also assess whether the network changes on the fast time scale within each window. We therefore evaluate the instantaneous rate of change of FC.

For each allegiance state, we plot the distribution of frame-to-frame FC variation, computed as the cosine distance (1 minus cosine similarity) between instantaneous FC matrices. At each time stamp *t*, the weighted FC is given by wavelet coherence between all region pairs. We then compute *v*(*t, t* + 1) ≡ *v*_*t*_, quantifying the change from *t* to *t*+1, and average these values over all consecutive time points within a window to obtain the windowed dFC speed ⟨*v*_*t*_⟩_*w*_ (Figure 2.b). This provides a fast-timescale measure aligned with the time resolution of the allegiance matrices used to construct the *dAm*.

From Figure 4.b (top row), the baseline allegiance state is not static: windows in this state show nonzero speeds, and—across recordings—baseline often exhibits the *highest* internal reconfiguration speed. Thus, links do change over time, yet do so without producing random module rearrangements, as indicated by the structured allegiance matrices. Across patients, baseline speed remains high, while allegiance structure can vary, meaning that baseline windows range from more gaseous” to more liquid” reconfiguration styles—but are always highly dynamic.

This contrast between weak FC and strong allegiance in the *Baseline* state is reversed in the *Seizure* state (Figure 4.b, second row). The state-averaged FC (left colormap) shows globally elevated off-diagonal values, reflecting widespread synchronization during seizure. The corresponding allegiance matrix still contains some coordinated groups—most visibly in the upper-left corner—but these are fewer and less pronounced. Whereas the baseline state exhibits a clear separation between region groups that consistently co-move (red blocks) and those that do not (blue stripes), this structure largely disappears during seizure, with blue dominating for this patient.

Thus, a fast but module-preserving reconfiguration at baseline is replaced by a slower, more module-mixing flow during seizure. The substantially reduced modularity in seizure windows (Figure 3) suggests that the weaker allegiance partly reflects the difficulty of extracting meaningful modules from networks that are, during seizure, only weakly modular. In other words, we could say that moving from baseline to seizure regimes, correspond to a “liquid-to-glass” transition.

The average FC of the third state (Figure 4.b, third row) shows a persistent block of strongly connected regions in the upper-left corner, linked to two smaller diagonal blocks. Its average allegiance matrix—defining this *Recovery* state—contains a few high-allegiance groups in the same upper-left portion, corresponding to the strongly connected FC regions. This part of the allegiance matrix resembles that of the *Seizure* state, whereas the remaining entries appear more “crystallized,” similar to the corresponding baseline pattern. The state therefore represents a transition between seizure and baseline in both connectivity and allegiance structure. This interpretation is supported by the *v*_*t*_ density plot, whose peak falls between those of the baseline and seizure states.

The fourth state exhibits FC and allegiance patterns distinct from the previous ones. In the FC matrix (Figure 4.b, bottom row), the 40Hz layer splits into two subgraphs—a smaller block in the upper-left corner and a larger one in the lower-right. The corresponding allegiance matrix (*MA*(*i, j*) colormap) shows the same division: two diagonal blocks whose nodes reconfigure in time in a coordinated manner within each subset but largely independently across subsets. This state thus features two “crystal-like” and mutually independent allegiance dynamics.

Notably, the two subnetworks correspond to temporal regions (top-left block) and frontal–parietal regions (bottom-right block). The associated *v*_*t*_ density plot shows a peak lower than in the preceding state, indicating a transient slowing of connectivity dynamics. This allegiance configuration differs substantially from the Baseline, Seizure, and initial Recovery states. We refer to it as a *“Recovery II”* state, noting that, depending on the recording, two or more such allegiance states may alternate throughout the extended post-ictal period.

The analyses in Figure 5.a provide a direct view of how the four allegiance states alternate over time. We plot the window-averaged dFC speed ⟨*v*_*t*_⟩_*w*_ and color each point according to its allegiance state. This representation confirms that the *Baseline* state exhibits the fastest connectivity dynamics (consistent with Figure 4.b), whereas the *Seizure* state shows the lowest ⟨*v*_*t*_⟩_*w*_, except for two pre-ictal transient epochs associated with clinician-annotated spike events and a brief perturbed interval of the 40Hz layer between 480 and 840 seconds.

**Figure 5.**
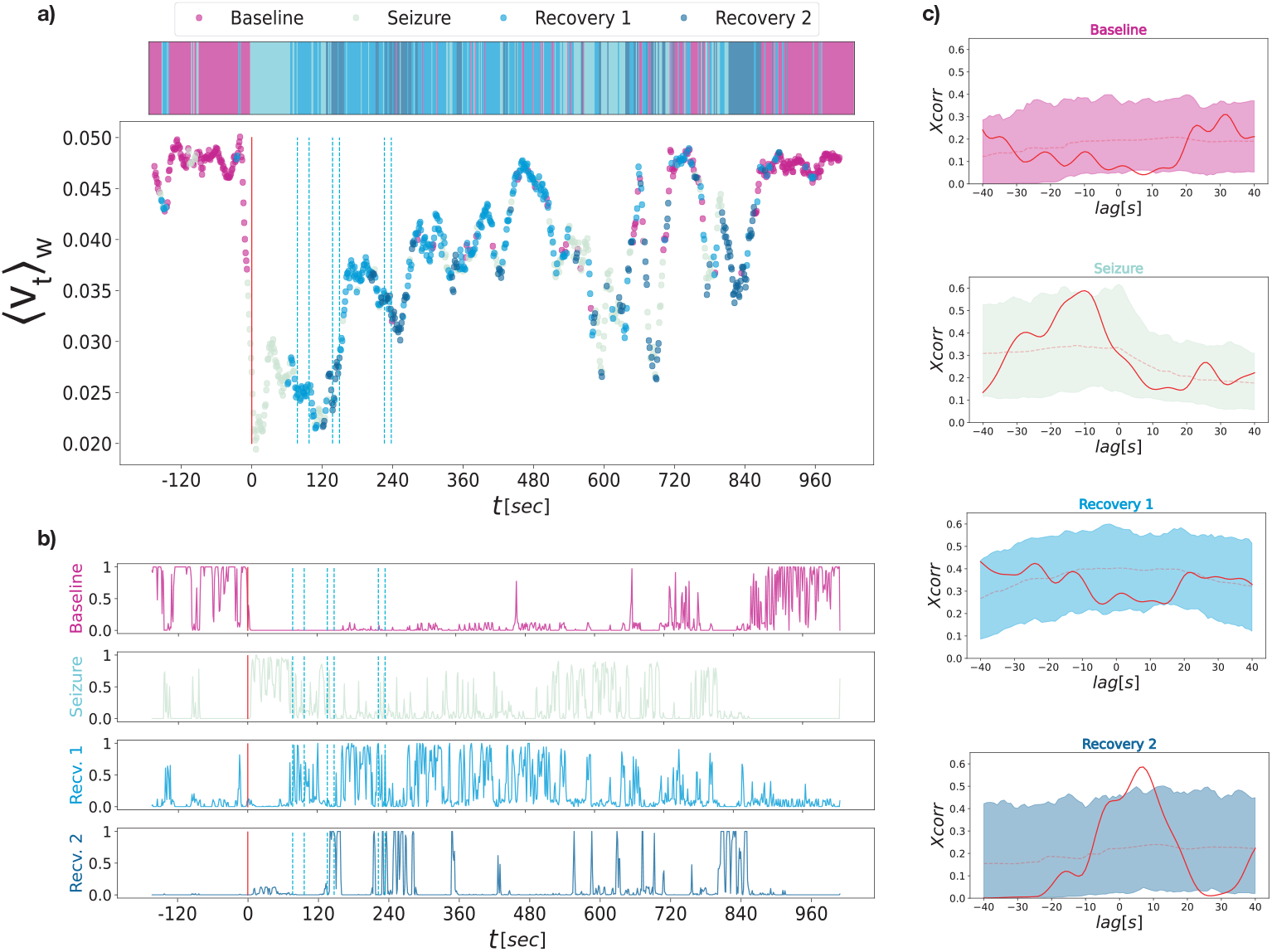
State-wise dFC-speed and symptom-relation: **a)** each dot in the plot corresponds to the average dFC-speed value within a time window, colored with the color of the corresponding allegiance state - here it is evident that the baseline state (magenta points at the begginning and end of the recording) is characterised by higher values of dFC-speed, whereas the seizure state has the lowest values, hinting at a sudden slowing down of the overall network dynamic reconfiguration; recovery 1 points are concentrated in periods of rising dFC-speed and on local maxima, whereas recovery 2 time windows happen around momentary slwoing downs of the dynamics and local minima. Each of the four curves in **b)** represent the probability *P* (*t*) ∈ [0, 1] of each time window *t* to belong to the baseline (magenta, top), seizure (light green, second plot), recovery-1 (light blue, 3*rd* plot) or recovery-2 (dark blue, bottom) states. Each of the four plots in **c)** represents the lagged cross-correlation between the peaks of the curves in **b)** of each state (baseline, seizure, recovery-1 and recovery-2) with the language dysfunction markers (blue lines); the shaded areas are comprised between the 5th and 95th percentiles of the values of cross-correlation computed for the peaks of *P* (*t*) of each state with a random permutation of the clinical markers in the [60*s*, 260*s*] interval.

The *Recovery* and *Recovery II* states alternate throughout the post-ictal period: the former appears during phases of increasing dFC-speed *v*_*t*_, while the latter accompanies sudden drops. Notably, these sharp decreases—marking entry into the Recovery II state—tend to occur near clinician-annotated transient language dysfunctions. This suggests that the fourth state may be systematically linked to aphasia episodes and that, speculatively, impaired information exchange during this slow, crystallized Recovery II regime could underlie the transient symptoms (see *Discussion*).

### 2.6 Allegiance state dynamics correlates with transient aphasia

The FC and 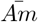 matrices in Figure 4.b represent archetypes of clusters associated with each allegiance state. However, each window reflects a distinct network reconfiguration with unique frames and sequences. To link allegiance dynamics with symptom occurrence, we adopt a soft classification approach, assigning each window not a single crisp label but a vector of similarities to all reference states. We implement this via a K-nearest-neighbour classifier, trained on window-specific dAm matrices labeled by spectral clustering (see *Methods*). The classifier outputs probabilities that may peak for specific states during strongly representative epochs, or assume intermediate values during smooth transitions or atypical windows. Systematic increases in the probability of a given allegiance state concurrent with language dysfunction indicate that the state is functionally related to the emergence of aphasia symptoms.

Figure 5.b shows the probabilities of each window belonging to the four allegiance states for the same recording as Figure 5.a. The magenta curve (top) represents the *Baseline* state, initially near 1 except for two brief drops, falling to 0 around seizure onset, and rising again in the final ∼ 4 minutes. The *Seizure* state (green) starts at 0, peaks during the Baseline drops (suggesting preictal spikes), and reaches its maximum between seizure onset and the first aphasia marker, before returning to 0 as Baseline recovers. The *Recovery* state rises near the first language dysfunction marker, fluctuates sharply until ∼ 540 seconds post-onset, then gradually declines to near-zero. The *Recovery II* state (dark blue, bottom) is rare but peaks near the last three clinical markers and intermittently until ∼ 860 seconds; it drops to 0 only when Baseline is fully restored.

Visual inspection suggests that drops in dFC speed *v*_*t*_ (Figure 5.a) co-occur with *Recovery II* peaks and aphasia symptoms. To quantify this, Figure 5.c shows lagged cross-correlation *XC*(*τ*) between allegiance-state assignment probabilities and symptom annotations, with significance assessed against a chance-level band from randomized symptom positions (see *Methods* for proper definition). Only the Seizure and *Recovery II* states show significant correlations, summarized by the *symptom relation index* Θ, which measures the area between *XC*(*τ*) and the 95% confidence interval. Here, Θ = 0.1256 for Seizure and Θ = 0.3231 for *Recovery II*, confirming the visual impression quantitatively.

[h!]

### 2.7 Interlayer co-allegiance

So far, we have analyzed each network layer independently. To study inter-layer relations, we compare module allegiance dynamics across frequency layers by computing the cosine similarity between allegiance matrices in the same time window across layers. This yields a time series of allegiance similarity for each layer pair, which we term the *multiplex inter-layer co-allegiance A*_*mpx*_ (Figure 6.a; see *Methods* for definition).

**Figure 6.**
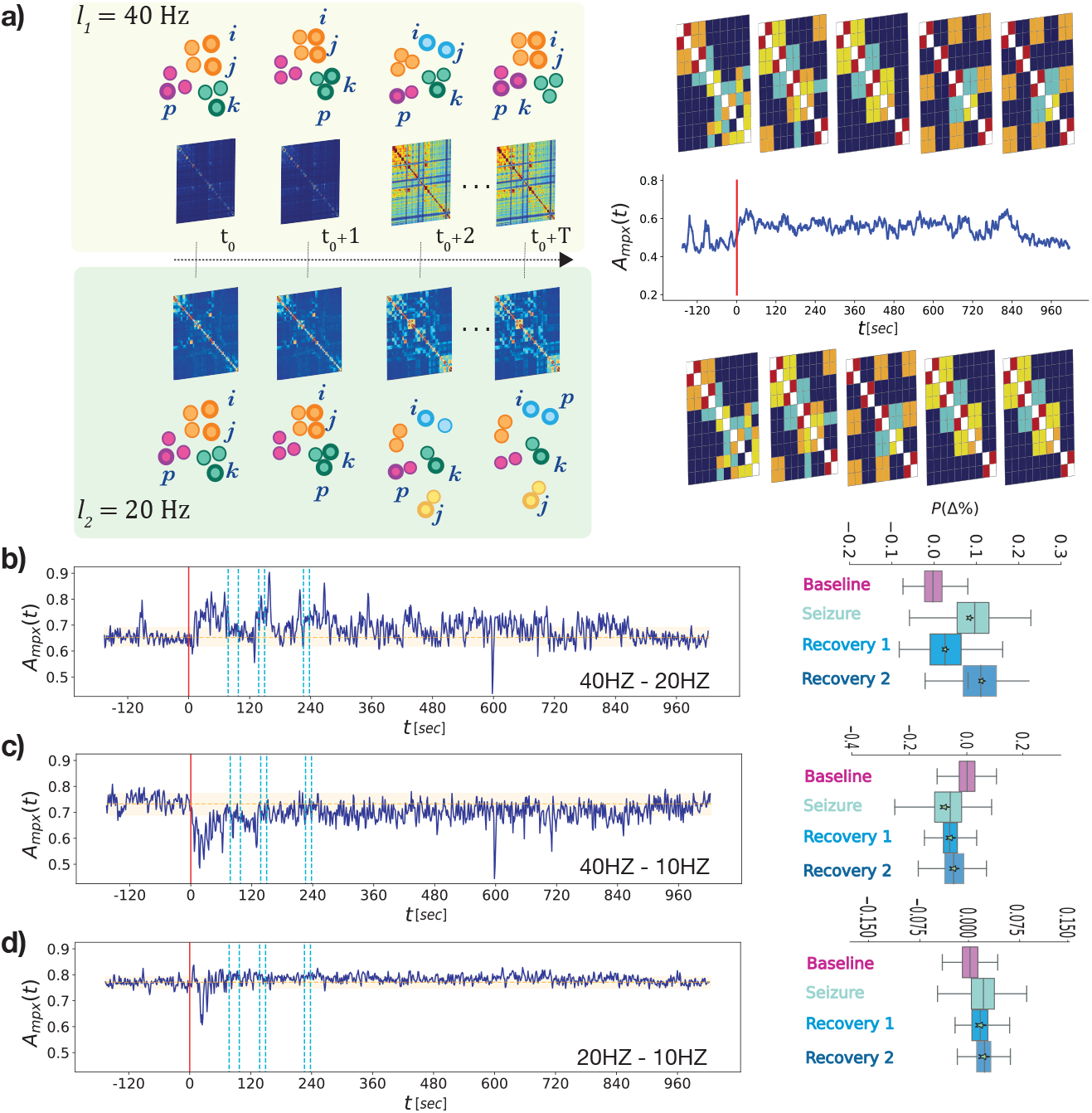
Inter-layer co-allegiance: **a)** For each pair of layers (*l*_1_ = 40Hz and *l*_2_ = 20Hz in the cartoon on the left) we compare the partition into modules at each time stamp by computing the cosine similarity of the relative module assignments - (right) the instantaneous inter-layer cosine similarity computed between the allegiance matrix of the 40Hz layer in a time window and the allegiance matrix of the 20Hz layer in the same window, recording 1. **b-c-d)** In each figure we plot: on the left, the curve of inter-layer co-allegiance (dark blue) and compare it to the average value of inter-layer co-allegiance found in the baseline state (orange dotted line, the orange-shaded area corresponds to values between the 5th and 95th percentile of baseline inter-layer co-allegiance values); On the right, each box describes the distribution of the interlayer co-allegiance within the corresponding allegiance state (baseline in magenta, seizure in turquoise, recovery 1 in light blue, recovery 2 in dark blue). The plots in **b)** correspond to the comparison of the 40Hz and 20Hz frequency bands of the representative recording, **c)** to 40Hz vs. 10Hz frequency bands and **d)** to 20Hz vs 10Hz, where the allegiance states have been retrieve by analysing the 20Hz layer.

We first examine the high-frequency layers at 40 and 20 Hz, which show the strongest seizure effects. Multiplex co-allegiance increases relative to baseline in all other states (Figure 6.b). Both Seizure and Recovery II states exhibit peaks beyond baseline variation. Percent changes relative to baseline (Figure 6.b, right) show positive shifts for all states, most pronounced in Seizure and Recovery II, indicating that 40Hz and 20Hz modules become more synchronized during seizure.

By contrast, 40Hz–10Hz co-allegiance drops during Seizure (Figure 6.c), with Seizure, Recovery, and Recovery II distributions shifted below baseline, reflecting decoupling that slowly re-establishes through recovery. For 20Hz–10Hz, co-allegiance also initially drops (Figure 6.d, left), but then increases for Recovery and Recovery II states, resembling 40–20Hz dynamics, though weaker.

In summary, network reorganization across layers is coordinated. At baseline, co-allegiance is weaker between 40Hz and 20Hz (*A*_*mpx*_ ≈ 0.65) and stronger with 10Hz (*A*_*mpx*_ ≈ 0.75). Seizure onset perturbs these patterns, enhancing fast-layer coupling and reducing 40–10Hz coupling, likely disrupting cross-frequency interactions required for language (see *Discussion*).

### 2.8 Comparison of different seizure recordings

After analyzing a reference recording, we compare results across seizures to identify similarities and variability (see *Discussion*). Figure 7 summarizes normalised descriptive features (non-normalised values can be found in Table 5) of allegiance states (columns) across recordings (rows) using radar plots showing:

- mean dFC-speed 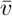;
- *symptom relation index* Θ;
- mean allegiance matrix 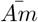;
- average 40–20Hz inter-layer co-allegiance.

**Figure 7.**
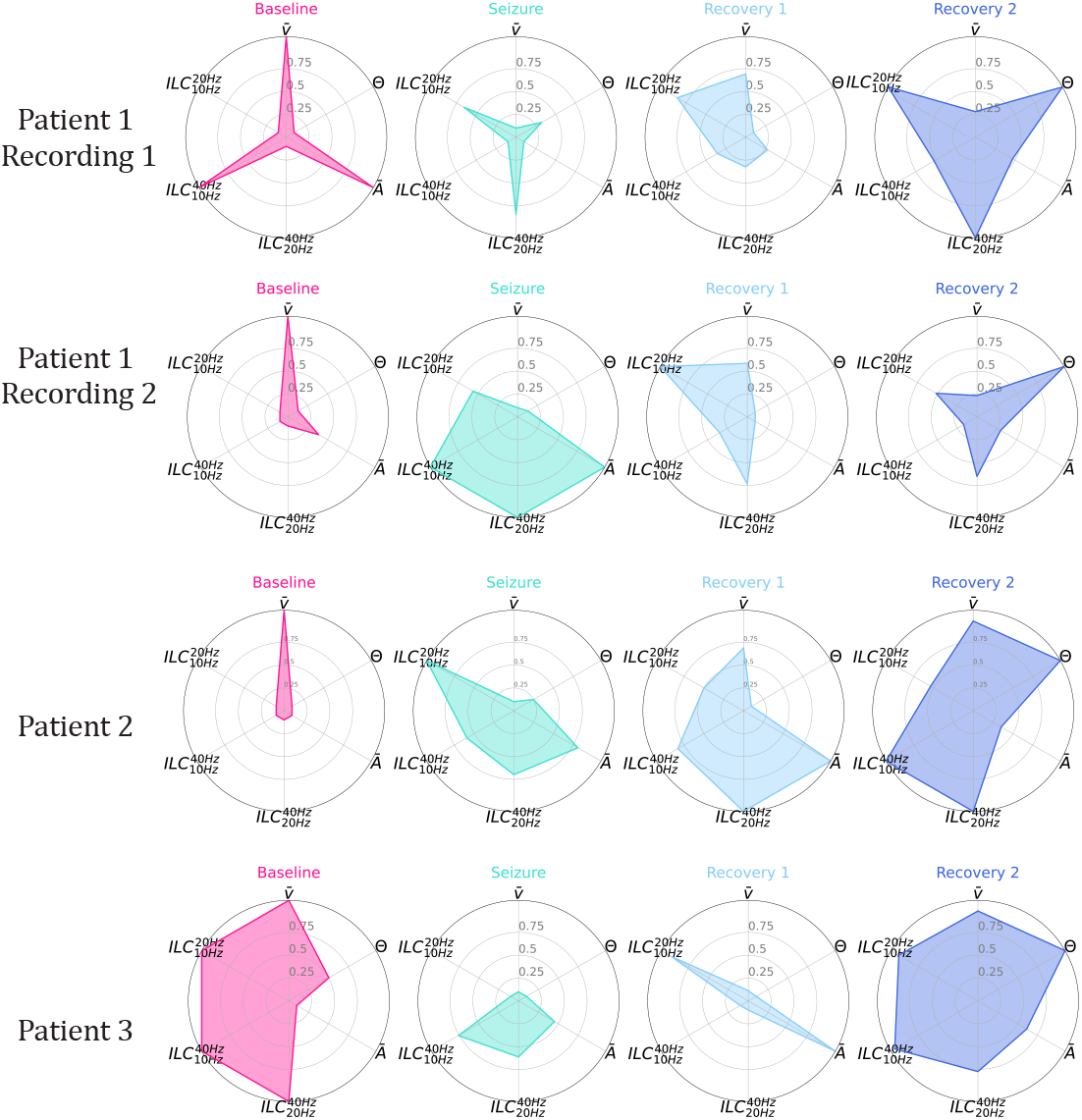
Allegiance states in the 40 Hz band: for each recording (two recordings of patient 1 and one recording each for patients 2 and 3) we show here four allegiance states by plotting on a radar plot four quantities: (from the top, clock-wise) the dFC-speed averaged on the time points of the corresponding state 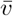; the Θ index of each state; the average over the allegiance matrix computed on the time points assigned to the relative state 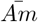; the average inter-layer co-allegiance computed over the corresponding allegiance states between the 40Hz and 20Hz layers. The four quantities that give rise to each of the four radar plots of a recording are normalised over the recording: a value of 1 of a quantity in a radar plot means maximum value of that quantity with respect to radar plots in the same row. The magenta radar-plots correspond to the baseline state; the light green plots correspond to seizure states, light and dark blue plots correspond to recovery 1 and 2, respectively.

Baseline consistently shows the highest 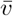, reflecting rapid reconfiguration, while Seizure has the slowest speed. Recovery states are intermediate, with Recovery I usually faster than Recovery II. Recovery II is most consistently distinguished by its strong correlation with language impairment (Θ).

Median allegiance strength 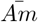 varies across states and recordings: in some seizures, baseline is highest and Seizure lowest; in others, the opposite occurs. Inter-layer 40–20Hz co-allegiance shows similar variability, usually increasing in Seizure and decreasing in Recovery, though some recordings reverse this pattern.

Overall, allegiance states display qualitative similarities across seizures, with baseline and Seizure dynamics markedly distinct and Recovery states transitional.Our metrics however were sensitive enough to capture subtle seizure- and patient-specific variations. Variability likely reflects heterogeneous baseline network configurations affecting seizure-induced dynamics (see *Discussion*).

## 3 Discussion

We introduced a framework for studying multi-frequency, time-varying data as a temporal multiplex network—a representation that has only recently begun to appear in computational work

[34] and remains largely absent in data-driven systems neuroscience. Our approach is motivated by the definition of a null model for the generation of synthetic temporal networks with fluctuating micro- and meso-scale (modules) structure, which reveals four prototypical reconfiguration styles. These states of temporal network dynamics motivate our empirical analysis and provide a conceptual baseline for interpreting real SEEG dynamics.

Although the present dataset is small (four recordings from three individuals), our approach builds on a substantial body of prior work highlighting the role of functional network dynamics in seizure evolution [20, 35–37]. Most previous studies, however, have relied on a single temporal resolution to characterise dynamic functional connectivity, typically focusing on comparisons between seizure-stage–specific representative frames or on global time courses of link dynamics.

Here, by explicitly combining two temporal resolutions, we characterise seizure stages as distinct states of unified network flow rather than bundles of stage-specific network configurations. This perspective helps reveal differences between phases such as the pre-ictal baseline and the late recovery period, in which the overall speed of network flow may already be similar and characteristic frames may appear comparable, yet allegiance remains disrupted, indicating a lower level of network order.

A further advance is that, unlike earlier studies restricted either to slow fMRI signals or to high-gamma power, our temporal multiplex representation captures coherence-based interactions across multiple frequency bands. Because oscillatory coherence supports information routing [38] and may organise neural representations across nested time scales [39–41], a multiplex description is particularly well suited to characterising flexible, multi-frequency communication.

This is especially relevant for language, where different frequencies pace the parsing of phonemes, syllables, words, and sentences [28, 29]. If dFC reflects ongoing information processing [1, 2, 42–44], then inter-frequency coordination should appear as intermediate (neither fully independent nor fully synchronous) alignment across layers. Consistently, baseline inter-layer co-allegiance values fall between zero and one (Figure 6b). At seizure onset, coupling between the 40Hz and 20Hz layers increases, suggesting disruption of hierarchical cross-frequency coordination important for language operations.

A key contribution of our framework is that it characterizes not only instantaneous network frames but also the flows through which they reorganize. Using a double time-scale, we describe fast reconfiguration within windows and slower evolution across windows. This leads to the notion of allegiance states, which summarize dynamic properties—such as reconfiguration speed and structured vs. random co-switching of nodes—rather than time-averaged connectivity alone [45]. Importantly, allegiance analysis bypasses the need to run full dynamic community detection [46–48] on very long, and high frequency temporal networks—an approach that is often computationally prohibitive in settings like ours—while still capturing the relevant cross-time modular structure. Speed proves particularly informative, allowing identification of similar allegiance states across seizures and patients [9, 10].

Contrary to interpretations of baseline as a stable configuration, our results show that baseline is characterized by the highest reconfiguration speed, indicating a flexible, continuously flowing network regime. Seizure onset induces a steep deceleration, consistent with the hyper-synchronization that constrains network variability, and recovery proceeds through intermediate states (Figure 4). Speed alone, however, cannot determine the style of reconfiguration: allegiance matrices specify which nodes move together and which fluctuate independently. For example, Patient 1’s baseline (Figure 7, top row) exhibits both high speed and structured allegiance, consistent with a rich, organized spatiotemporal flow that may boost computational capacity [49]. In seizures, high allegiance often reflects frozen connectivity or noise correlations rather than functional organization.

Baseline allegiance strength varies across recordings (Table 5), likely due to differences in cognitive state, seizure type [50, 51], and baseline duration. Group-level analyses could be performed to harness variance and better highlight common aspects [52–54]. We chose here however on purpose to focus on single seizure-level analyses to highlight the capability of our method to capture subtle within- and between-patient differences, supporting personalized neuromarker development.

Importantly, the recovery phase is characterized by two states. A first recovery state gradually restores high-speed reconfiguration, while a second, transient “Recovery II” state slows dynamics again. In “Recovery II,” allegiance matrices consistently segregate into temporal versus fronto-parietal subnetworks (Tables 2,3,4). This functional decoupling aligns with regions necessary for language function and explains the elevated Θ values and association with aphasia. Notably, this “solid-like” state parallels one of the null model’s four reconfiguration styles, underscoring that these dynamical regimes are not idiosyncratic but reflect general organizational constraints.

**Table 2:**
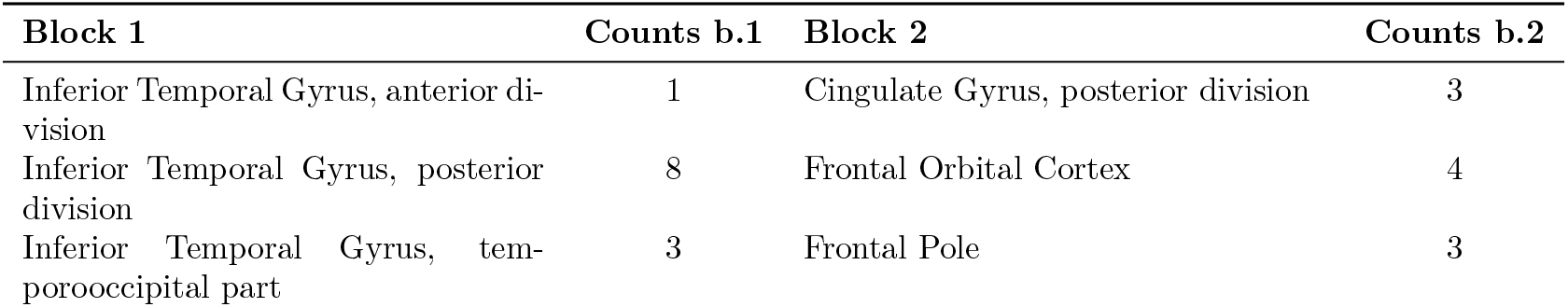
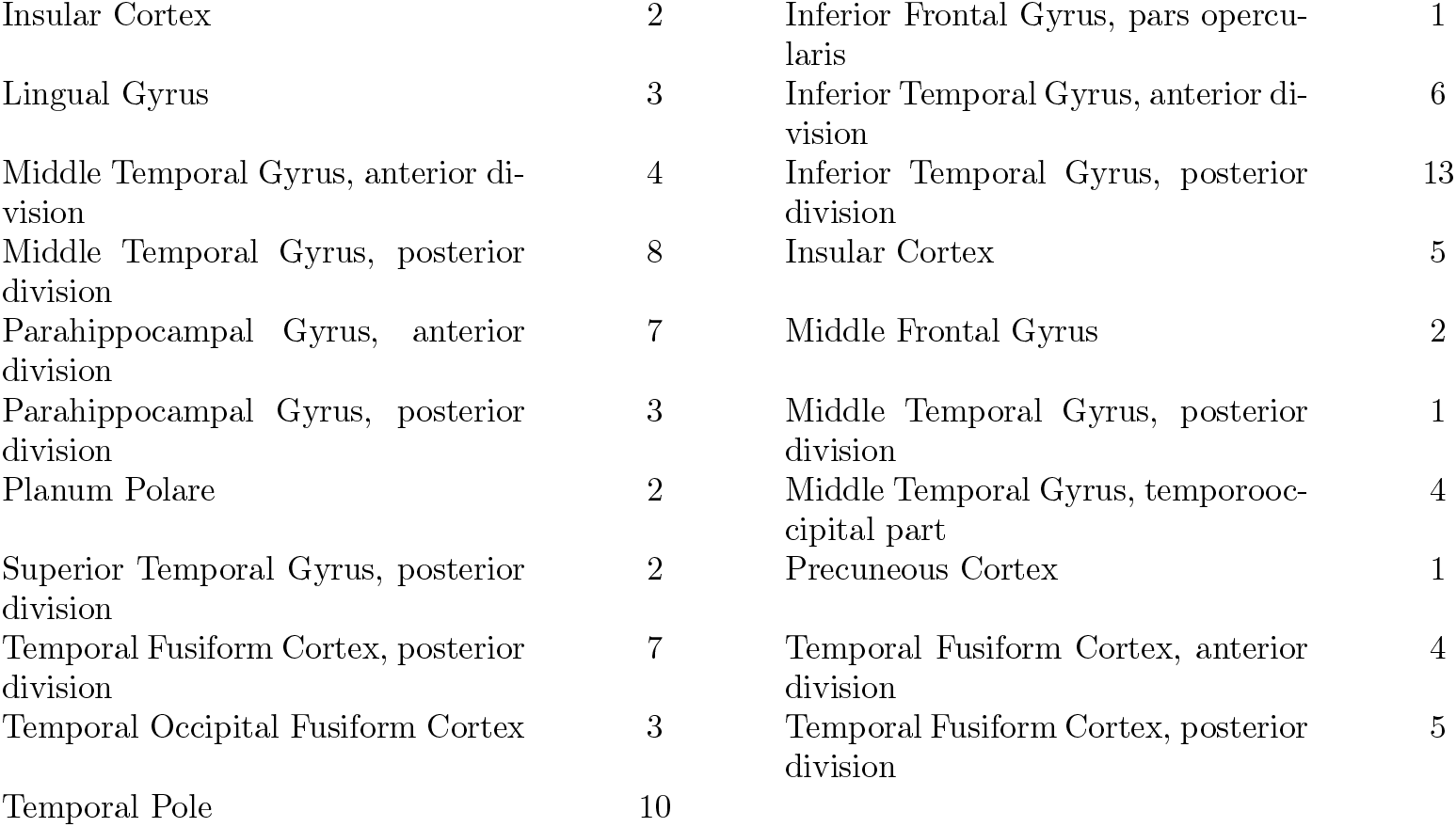
Brain areas per block during Recovery 2 state, Patient 1.

**Table 3:** Brain areas per block during Recovery 2 state, Patient 2.

| Block 1 | Counts b.1 | Block 2 | Counts b.2 |
| --- | --- | --- | --- |
| Inferior Temporal Gyrus, posterior division | 5 | Frontal Operculum Cortex | 5 |
| Middle Temporal Gyrus, anterior division | 10 | Frontal Orbital Cortex | 5 |
| Middle Temporal Gyrus, posterior division | 16 | Frontal Pole | 34 |
| Parahippocampal Gyrus, posterior division | 1 | Inferior Frontal Gyrus, pars opercularis | 6 |
| Planum Polare | 2 | Inferior Frontal Gyrus, pars triangularis | 2 |
| Superior Temporal Gyrus, anterior division | 3 | Insular Cortex | 1 |
| Temporal Fusiform Cortex, posterior division | 4 | Middle Frontal Gyrus | 5 |
| Temporal Pole | 17 | Middle Temporal Gyrus, anterior division | 7 |
|  |  | Parahippocampal Gyrus, anterior division | 5 |
|  |  | Parahippocampal Gyrus, posterior division | 1 |
|  |  | Temporal Fusiform Cortex, anterior division | 1 |
|  |  | Temporal Fusiform Cortex, posterior division | 2 |

**Table 4:** Brain areas per block during Recovery 2 state, Patient 3.

| Block 1 | Counts b.1 | Block 2 | Counts b.2 |
| --- | --- | --- | --- |
| Heschl’s Gyrus (includes H1 and H2) | 4 | Inferior Temporal Gyrus, temporooccipital part | 2 |
| Insular Cortex | 10 | Lingual Gyrus | 3 |
| Parahippocampal Gyrus, anterior division | 4 | Middle Temporal Gyrus, temporooccipital part | 5 |
| Planum Polare | 4 | Temporal Occipital Fusiform Cortex | 1 |
| Planum Temporale | 1 |  |  |
| Superior Temporal Gyrus, anterior division | 2 |  |  |

**Table 5:** Absolute values of the features displayed in Figure 7 for all recordings.

| Recording | State | $\bar{v}$ | $\Theta$ | $\bar{A}$ | $ILC_{20Hz}^{40Hz}$ | $ILC_{10Hz}^{40Hz}$ | $ILC_{10Hz}^{20Hz}$ |
| --- | --- | --- | --- | --- | --- | --- | --- |
| Patient 1 rec. 1 | Baseline | 0.048427 | 0.000000 | 0.269140 | 0.652322 | 0.732896 | 0.771437 |
|  | Seizure | 0.029320 | 0.130109 | 0.075238 | 0.706959 | 0.676124 | 0.780425 |
|  | Recovery 1 | 0.040629 | 0.000000 | 0.109371 | 0.668743 | 0.690736 | 0.783780 |
|  | Recovery 2 | 0.032766 | 0.579639 | 0.147182 | 0.725582 | 0.700006 | 0.787612 |
| Patient 1 rec. 2 | Baseline | 0.003530 | 0.000000 | 0.221519 | 0.702906 | 0.766885 | 0.822679 |
|  | Seizure | 0.000779 | 0.000000 | 0.348581 | 0.812069 | 0.886870 | 0.825763 |
|  | Recovery 1 | 0.002115 | -0.005870 | 0.170053 | 0.772118 | 0.796673 | 0.829347 |
|  | Recovery 2 | 0.001146 | 0.189989 | 0.205292 | 0.763039 | 0.774996 | 0.825457 |
| Patient 2 | Baseline | 0.027833 | 0.000000 | 0.061227 | 0.805935 | 0.684796 | 0.811552 |
|  | Seizure | 0.016485 | 0.134928 | 0.167022 | 0.828347 | 0.741896 | 0.850336 |
|  | Recovery 1 | 0.023148 | 0.000000 | 0.210452 | 0.843399 | 0.769216 | 0.827033 |
|  | Recovery 2 | 0.026493 | 0.907741 | 0.099023 | 0.843395 | 0.800636 | 0.828520 |
| Patient 3 | Baseline | 0.035345 | 0.096972 | 0.085706 | 0.756283 | 0.685543 | 0.844199 |
|  | Seizure | 0.027083 | 0.000000 | 0.098120 | 0.728054 | 0.670637 | 0.771065 |
|  | Recovery 1 | 0.030225 | 0.000000 | 0.120874 | 0.698316 | 0.641979 | 0.834501 |
|  | Recovery 2 | 0.029053 | 0.239141 | 0.103834 | 0.737310 | 0.683400 | 0.836820 |

Seizure effects are strongest in the 40Hz layer, with 20Hz showing similar but weaker patterns; slower frequencies appear less affected. Unipolar recordings enhance detection of these dynamics compared to bipolar ones, likely because they better capture distributed collective activity rather than noise [55–59]. Methodological constraints may underlie the muted effects in slower bands, suggesting room for layer-specific optimization in future work.

While our analyses emphasize temporal organization over spatial detail, global features such as modularity and strength already reveal functionally meaningful changes. Incorporating meta-connectivity or edge-centric analyses [10, 60] could enhance spatial specificity once larger datasets become available.

In summary, we demonstrate that network dynamics themselves—captured by dFC-speed and modular reconfiguration style—constitute meaningful observables for neural function. Each allegiance state corresponds to a distinct mode of exploring FC configurations: fast and flexible at baseline, viscous and constrained during seizure, and selectively slowed or reorganized during recovery. Critically, two such states align with clinician-annotated language impairments, suggesting that dynamic reconfiguration styles may offer novel biomarkers for both healthy cognition and its disruption in epilepsy.

## 4 Methods

### 4.1 Null model

Our proposed model represents a multilevel, or hierarchical variation of the Dynamic Stochastic Block Model (DSBM), a benchmark generative model for synthetic temporal networks. In our model, we consider *N* nodes, assigned to *n*_1_ first-level modules of the same size (*N/n*_1_). All nodes in the same first-level modules are then also assigned to one of the *n*_2_ *< n*_1_, randomly selected, second-level, meso-scale modules. Similarly to the regular static SBM, nodes belonging to the same second-level module are connected by a link with probability *p*_*in*_, while nodes assigned to two different second-level modules are connected with probability *p*_*out*_ *< p*_*in*_. The static graph generated via this procedure thus represent the initial snapshot of the temporal network. At each discrete time step *t*, in fact, the evolution of the synthetically generated network is influenced by two competing stochastic processes:

- a first process affects all edges in the network: each edge in the network is resampled according to the original parameters of the model with probability *p*_*change*_, while it is preserved with *link-persistency* probability 1 − *p*_*change*_;
- a second process aims at stabilizing the first-level modules: all nodes in each first-level module are assigned to a new second-level module with probability *p*_*switch*_, or see no change in their second-level module assignment with probability 1 − *p*_*switch*_. If a first-level module (all of its nodes) has been assigned to a new second-level module, all of the edges connected to its nodes are rearranged in order to achieve a density of connections with nodes assigned to the same, new second-level community, w.r.t. the rest of the network that is equal to *p*_*in*_*/p*_*out*_.

The free parameters of our model are therefore: *p*_*in*_, *p*_*out*_, *p*_*change*_, *p*_*switch*_.

### 4.2 Recordings

We analysed four peri-ictal SEEG recordings from three patients with pharmacoresistant temporal lobe epilepsy who exhibited transient post-ictal aphasia. Each patient underwent clinical implantation for presurgical evaluation; electrode sampling varied according to clinical needs and covered temporal, frontal and insular regions to different extents. Tables 1-4 summarise the number of implanted contacts, anatomical coverage, recording durations and seizure counts for each patient. The SEEG recordings were performed using intracerebral multiple contact electrodes (10 to 15 contacts, length: 2 mm, diameter: 0.8 mm, 1.5 mm apart from edge to edge) implanted according to Talairach’s stereotactic method [61]. The strategy of electrode positioning was established in each patient based upon hypotheses concerning the localization of the epileptogenic zone (EZ), with the aim of defining subsequent cortectomy. These hypotheses of the likely EZ localization were based on noninvasive presurgical assessment including detailed clinical history, surface video-electroencephalographic (EEG) recording, MRI, and 18FDG-PET scanner. Electrode positions were therefore not standardized across the present series, but were chosen according to the individual clinical characteristics of each case. However, electrode coverage in all cases included at least several cortical contacts being parts of the language network: i.e., temporal lateral and basal temporal region, inferior frontal gyrus, anterior part of the insula [62]. Implantations were unilateral or bilateral depending on the individual features of each case. Signals were recorded using a 256-channel Deltamed system, at a 1024-Hz sampling frequency. We emphasise that the dataset is small, heterogeneous and clinically driven; consequently, our goal is not to perform population-level inference but to demonstrate a methodological framework that can robustly characterise dynamic connectivity in single-patient intracranial data.

### 4.3 Wavelet coherence

In order to compute the wavelet coherence between each pair of signals, and thus construct a graph representation of the data, we decompose signals in the time-frequency domain along the Morlet wavelet family. The Morlet wavelet is defined for frequency *f* and time *τ* as:

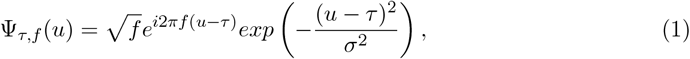

so that Ψ_*τ,f*_ (*u*) is the product of a sinusoidal wave at frequency *f* with a Gaussian function centered at time *τ*, with a standard deviation *σ* proportional to 1*/f* . The wavelet transform of a signal *x*(*u*) is a function of time *τ* and frequency *f* given by the convolution with the Morlet wavelet:

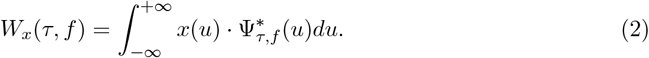

From the wavelet transform of two signals *x* and *y*, we can define the wavelet cross-spectrum between *x* and *y* around time *t* and frequency *f* :

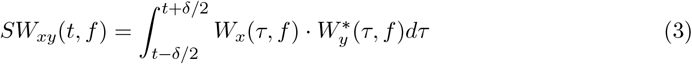

where *δ* is a scalar that depends on the frequency. The wavelet coherence *W.C*.(*t, f*) is defined at time *t* and frequency *f* by:

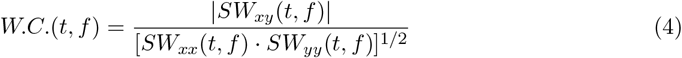

where *W.C*.(*t, f*) has values between 0 and 1. We compute the wavelet coherence *W.C*(*t, f*) for each pair of signals in 5 frequency bands peaked around 1Hz, 5Hz, 10Hz, 20Hz and 40Hz, where the width of the time integration *δ* = 6*/f*, and thus the width of each of the frequency bands equals 1*/*6 of the peak frequency. We define a significance threshold for the coherence values by computing the coherence for pairs of randomly generated oscillating signals (500 iterations), and thus consider as significant values measured on empirical data only those that lie above the 95% confidence interval of the distribution of coherence of the random data. The sampling frequency of the recording is 1024Hz, and we down-sample the time series of the coherence between each pair of signals by a factor 12, with a resulting time resolution of each layer of the temporal-multiplex of 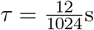.

### 4.4 Dynamic allegiance states

Several approaches have been developed to detect the presence of densely interconnected communities in temporal networks. In this work, given the size and complexity of the data we analyse, we focus, rather then on retrieving purely *temporal* communities, on investigating the mutual relations among nodes with respect to their instantaneous (static, computed at each time stamp) community memberships. In particular, we retrieve the instantaneous partition of the network on each of its snapshots and then compute, within a desired time window, the probability of each pair of nodes to be found in the same community at the same time.

#### 4.4.1 Allegiance

We do so detecting the instantaneous community partitions by a modularity based approach (the Louvain algorithm, [63]). We set the resolution parameter of the Louvain algorithm to *γ* = 1, knowing that such value favors the detection of larger communities. For each time point we ran Louvain 200 times thus obtaining 200 partitions. For each partition, we compute another definition of Modularity, introduced in [64], where the sampled network is compared to a different null model, instead of the classic configuration model. In fact, for each partition P we compute the modified modularity:

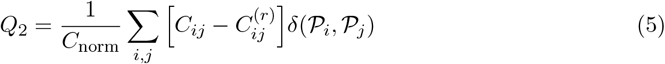

where *C* is the empirical instantaneous wavelet coherence matrix at a given time, *C*_*norm*_ = ∑_*i,j*_ *C*_*ij*_ is a normalization factor, 𝒫_*i*_ and 𝒫_*j*_ are the community assignments of nodes *i* and *j* in partition 𝒫, and *C*^(^*r*) corresponds to the *random* component of the empirical matrix *C* as defined in equation 13 in [64]. Such null model, rooted in Random Matrix Theory is referred to as the noise-only null model *ρ*^*MG*2^ in [65]. We thus retain as optimal partition among the 200 obtained via Louvain, the partition maximizing *Q*_2_. Within a time window *w* of 200 time points (∼2.34 seconds), we compute the so called *module-allegiance*, or *association* matrix [66] 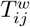, whose matrix element corresponds to the probability

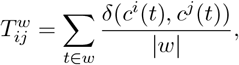

where |*w*| is the length of the time window *w, c*^*i*^(*t*) and *c*^*j*^(*t*) are the community assignments of nodes *i* and *j* at time *t*, respectively, and the Kronecker delta *δ*(*c*^*i*^(*t*), *c*^*j*^(*t*)) therefore equals 1 when the two nodes are assigned to the same community. We then slide the time window by a lag *τ* = 100 pts. (*τ* = 1.171 seconds), and compute the allegiance in the next window. Consecutive windows have therefore a 50% overlap in time.

#### 4.4.2 Dynamic Allegiance

The layer-wise Dynamic Allegiance, *dAm*^*l*^, shown in Figure 4 for the 40Hz frequency band, is obtained by computing the cosine similarity between the upper triangular part (linearized into an *N* * *N* − 1*/*2 vector *V* ^*l*^(*t*)) each allegiance matrix computed in the window *t*^*^ ∈ [*L*_*w*_, *L*_*w*_ + *τ*, …, *T/τ*] (where *τ* is the 100 time points lag by which we slide the time window) and all of the other allegiance matrices,i.e. :

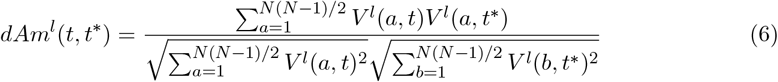

The matrix element *dAm*^*l*^(*t, t*^*^) therefore ranges from 0, when the two allegiance matrices in windows *t* and *t*^*^ are completely different, to 1, when the two are exactly identical. The blockwise structure of a layer’s *dAm* suggests the existence of epochs, or states, in which the allegiance matrices computed in the corresponding time windows are similar: these are states in which the allegiance dynamics of nodes (brain regions) is in a stable and persistent regime, that we refer to as *allegiance states*.

### 4.5 Clustering and KNN classification

#### 4.5.1 Clustering on the *dAm*: allegiance states

In order to extract periods of time (clusters of time windows) characterized by similar module allegiance of nodes, we use a spectral clustering method (implemented in the scikit-learn python library [67]) where the *dAm* matrix serves as the distance metric between time-points. Since the method requires an a-priori choice of the number of clusters, we iterate the clustering for different numbers of clusters, thus selecting the result yielding the maximal silhouette score. The silhouette score of a partition into clusters of a dataset is equal to 1 (maximum) when there is 0 overlap between clusters and all data points have been assigned to the right cluster; it is 0 when there are overlapping clusters and −1 when all data points have been assigned to the wrong cluster. The number of allegiance states is not fixed a priori and varies across recordings. In most cases, four clusters are identified, while in some frequency bands (multiplex network layers) of some recordings an additional recovery-related cluster was retrieved. This cluster accounted for a small fraction of windows and displayed heterogeneous dynamical properties, consistent with a transitional or mixed state. Given its limited temporal weight and lack of consistent structure, it was not assigned a specific functional interpretation.

#### 4.5.2 Unsupervised classification for soft allegiance-state labeling of time-windows

We use the K-nearest-neighbor unsupervised classification method (scikit-learn implementation), trained on the allegiance state labels of all the time windows, to obtain a soft classification of each the latter. This allows us to represent each module allegiance matrix computed in the relative time-window as a (in the case of **Patient 1**, as presented in Figure 5.b) 4-dimensional normalized vector: each component of this vector represents the probability of the time-window to belong to the corresponding allegiance state (Baseline, Seizure, Recovery 1, Recovery 2).

### 4.6 Symptom relation index Θ

To quantify the association between allegiance-state expression and clinician-annotated symptoms, we compute the lagged cross-correlation *XC*(*τ*) between the time series of allegiance-state assignment probabilities and the binary symptom time series (Figure 5c). Significance is assessed relative to a surrogate distribution obtained by randomising symptom timings.

#### Surrogate distribution

We generate *S* = 1000 surrogate symptom sequences 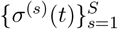 by circularly shifting the original symptom annotation by a random lag. For each surrogate we compute its cross-correlation with the allegiance-state probabilities, yielding a set of surrogate curves

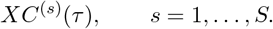

At each lag *τ*, the empirical surrogate distribution 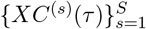 defines a null model. The 95% chance band shown in Figure 5c corresponds to the pointwise 2.5–97.5 percentile interval of this distribution.

#### Symptom relation index

We define the *symptom relation index* Θ as the integrated excess of the empirical cross-correlation above the upper 95% confidence bound of the surrogate distribution:

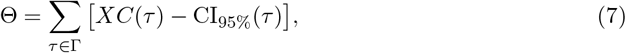

where the set of significant lags Γ Is

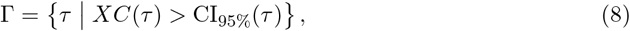

and CI_95%_(*τ*) denotes the 95th percentile of the surrogate distribution at lag *τ* . Thus, Θ quantifies the area by which the observed cross-correlation exceeds chance expectations across all significant lags *τ* .

### 4.7 dFC-speed

The instantaneous dFC-speed *v*_*t*_ as visually introduced in Figure 1.b (top) is defined as the cosine similarity between the FC matrix at time *t* and that corresponding to the previous, adjacent time point *t* − 1 (linearized into *N* * *N* − 1*/*2 vectors *FC*_*vec*_(*t*), *FC*_*vec*_(*t* − 1)):

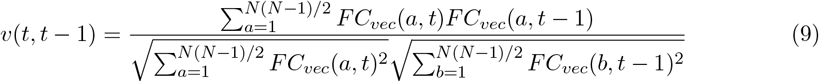

while the time-window-averaged dFC-speed ⟨*v*_*t*_⟩_*W*_ corresponds to the average value of the instantaneous dFC-speed *v*_*t*_ within a 200 time points (∼ 2.34 seconds) time-window *W* (the same windows for each of which the module allegiance was computed).

### 4.8 Interlayer co-allegiance

We define the interlayer co-allegiance between each pair of layers (*l, l*^*^) as the cosine similarity between the allegiance matrix computed in a time window in layer *l* (specifically, the upper triangular part of the matrix, linearized into an *N* * *N* − 1*/*2 vector *V* ^*l*^(*t*)), with the allegiance matrix computed in the same window layer *l*^*\**^, for all the time windows:

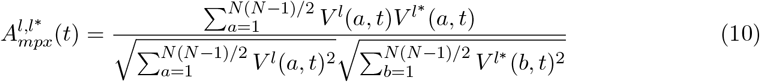

## Data availability statement

The conditions of our ethics approval do not permit public archiving of anonymised study data. Readers seeking access to the data should contact Dr. Agnès Trebuchon. Access will be granted to named individuals in accordance with ethical procedures governing the reuse of clinical data, including completion of a formal data sharing agreement.

