## Supplementary Info for "States of dynamic connectivity flow in temporal multiplex networks: a case study in human epilepsy and postictal aphasia"

### **Supplementary Information:** States of functional connectivity flow and their multiplex dynamics in human epilepsy and postictal aphasia

Nicola Pedreschi[1,2,3,⋆], Agnès Trebuchon[2,4], Alain Barrat[1], and Demian Battaglia[2,5]

**1 Aix-Marseille Univ, Université de Toulon, CNRS, CPT, Turing Center for Living Systems, Marseille, France**

**2 Aix-Marseille Université, Inserm, INS, Institut de Neurosciences des Systémes, Marseille, France**

**3 Mathematical Institute, University of Oxford, Oxford, UK**

**4 Assistance Publique Hôpitaux de Marseille (APHM), Service d'épileptologie, Marseille, France**

**5 University of Strasbourg Institute for Advanced Studies (USIAS), Strasbourg, France**

⋆ ****

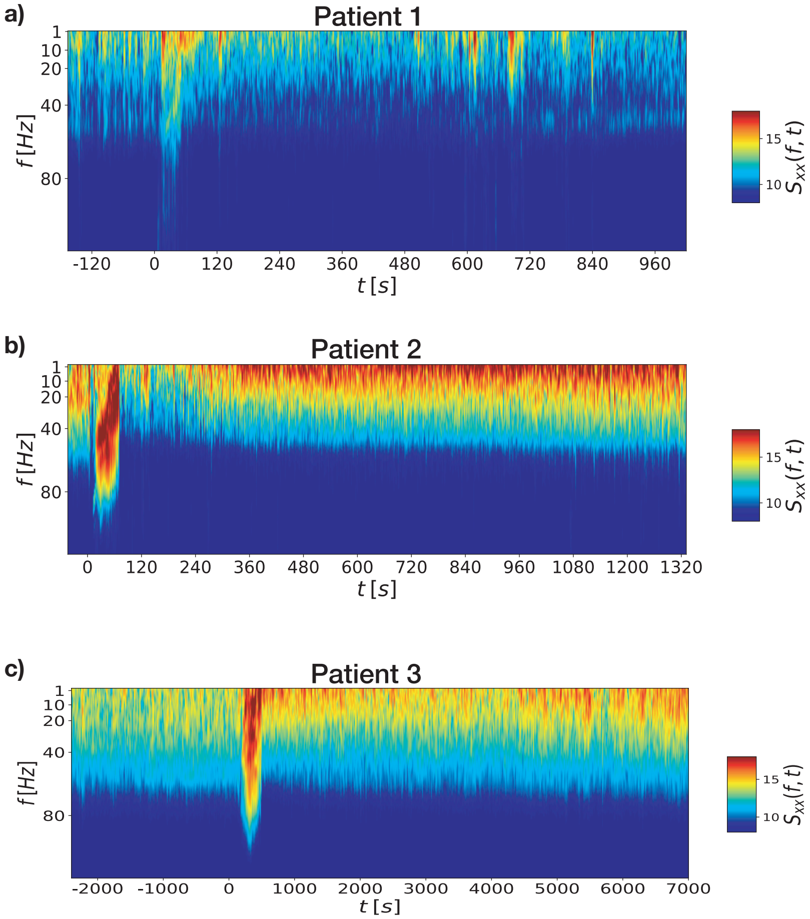


**Figure 1.** Power spectrum in the time-frequency domain of the mono-polar time series of an individual contact, for each of the three patients. On the x-axis is time, in seconds: the 0 on the x-axis corresponds to the seizure onset.

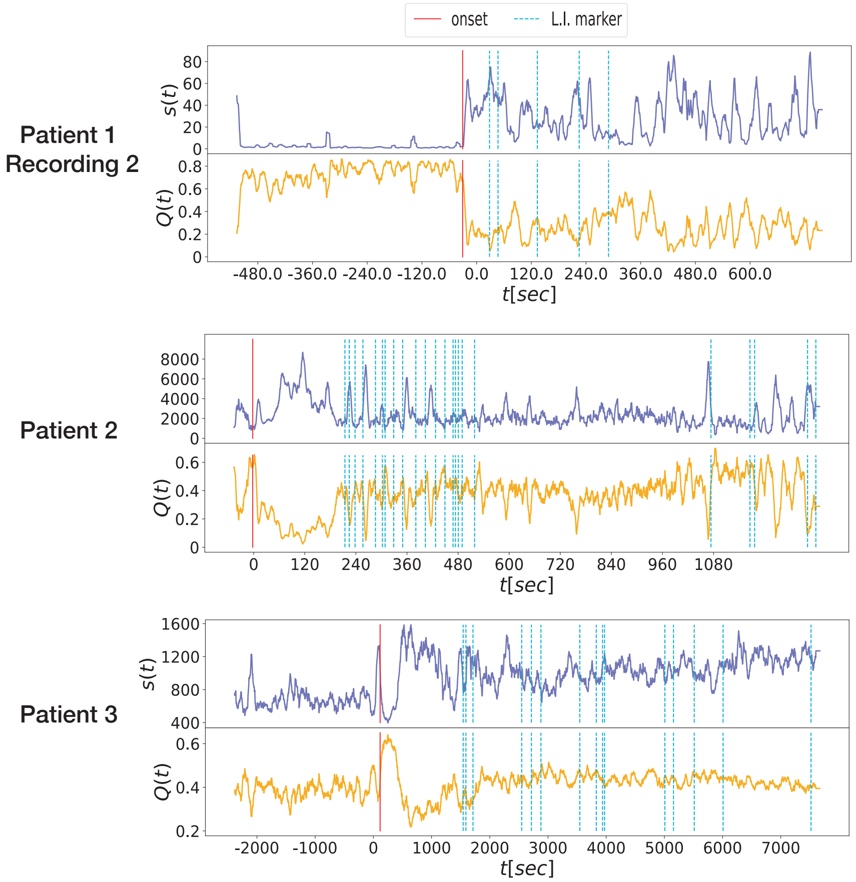


**Figure 2.** Time series of strength $s(t)$ and modularity $Q(t)$ for the $40Hz$ layer of each of the 3 recordings whose analysis are not displayed in the main text. The zero on the x-axis corresponds to seizure onset (red, vertical line) whereas the light-blue dotted lines correspond to the language impairment markers provided by the clinicians.

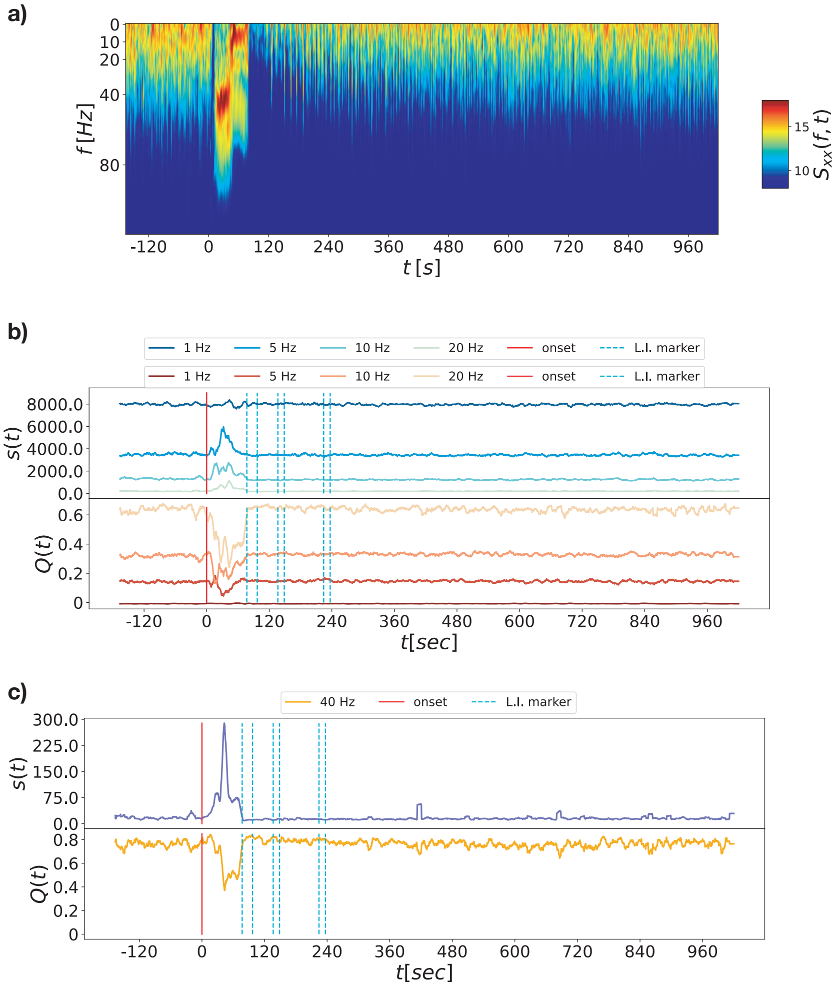


**Figure 3.** **a)**: power spectrum in the time-frequency domain of the bipolar time series of an individual contact for the first recording of Patient 1 (bipolar version of 1.a). **b-c)** correspond to the same analysis presented in **Figure 3** in the main text, but performed on the bipolar time-series: **b)** the curves in the top (bottom) plot correspond to the time series of instantaneous total strength $s(t)$ (modularity $Q(t)$) of four frequency bands, namely, the 1Hz frequency band in dark blue (dark red), 5Hz in blue (red), 10 Hz in light blue (dark orange), 20Hz in light-green (light-orange). In **c)** are the curve of $s(t)$ for the 40Hz frequency band in the top plot, blue-navy line, and of the modularity $Q(t)$ in the bottom plot, orange line. The vertical lines in the plots represent the clinical markers: the seizure onset in red, and the language dysfunction markers are represented by the cyan dotted lines.

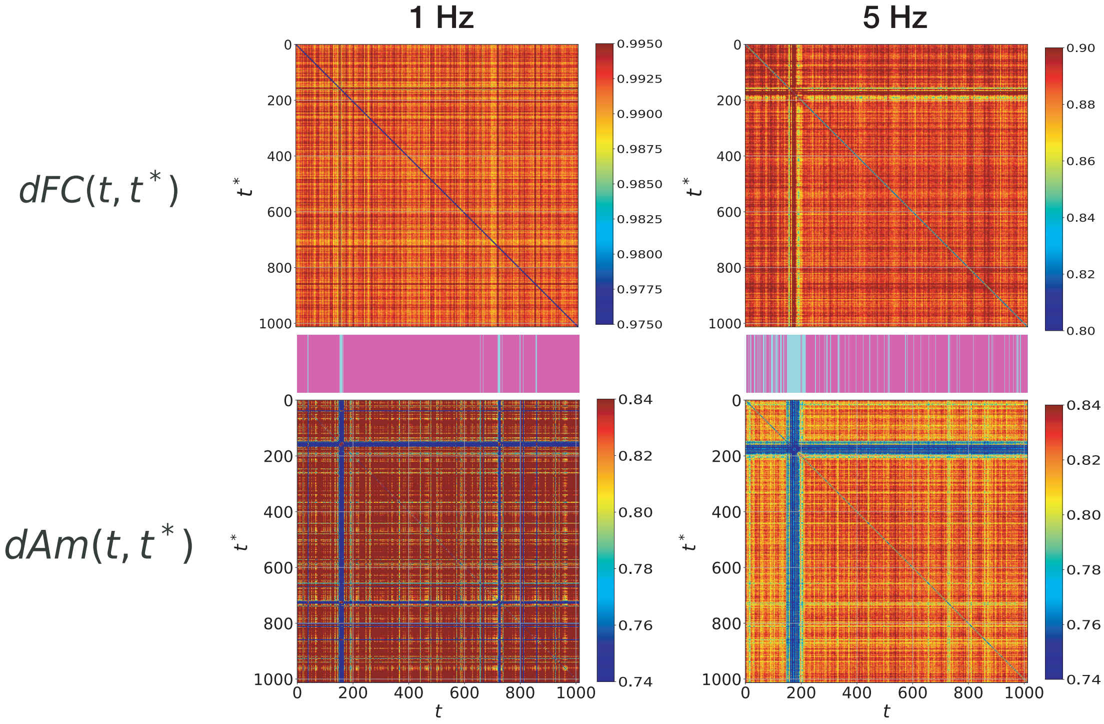


**Figure 4.** The **dFC** and **dAm**, as presented in **Figure 4.a** in the main text, computed for the 1Hz and the 5Hz layers. The barcode in between the two matrices, in each layer, represents the sequence of allegiance states: in these two layers, only two states where retrieved, namely a baseline state (magenta bars in the barcode) and a seizure state (aqumarine).

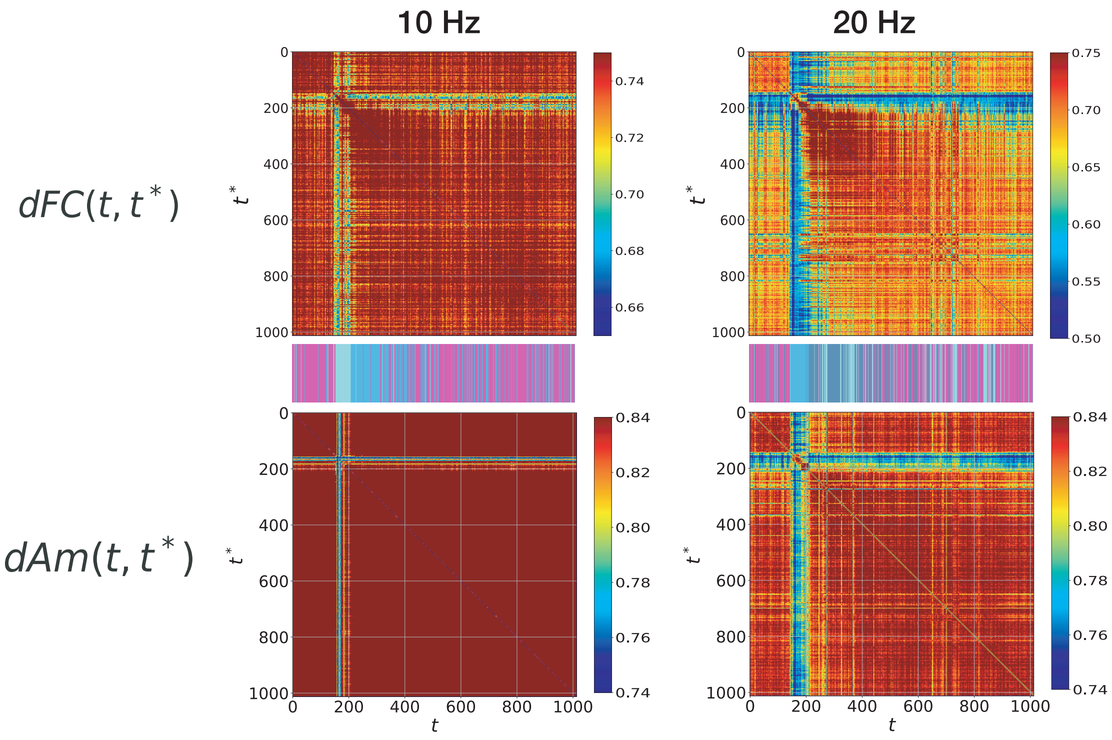


**Figure 5.** The **dFC** and **dAm**, as presented in **Figure 4.a** in the main text, computed for the 10Hz and the 20Hz layers of the first recording of Patient 1. The barcode in between the two matrices, in each layer, represents the sequence of allegiance states. In these two layers more than two states where found: the 10Hz layer has three allegiance states, namely, a baseline state (magenta) a seizure state (aqumarine) and a recovery state (light blue); the 20Hz layer undergoes four states, as the 40Hz layer presented in the main text, and has a baseline state (magenta),seizure state (aquamarine), a recovery 1 (light blue) and a recovery 2 (dark blue) states.

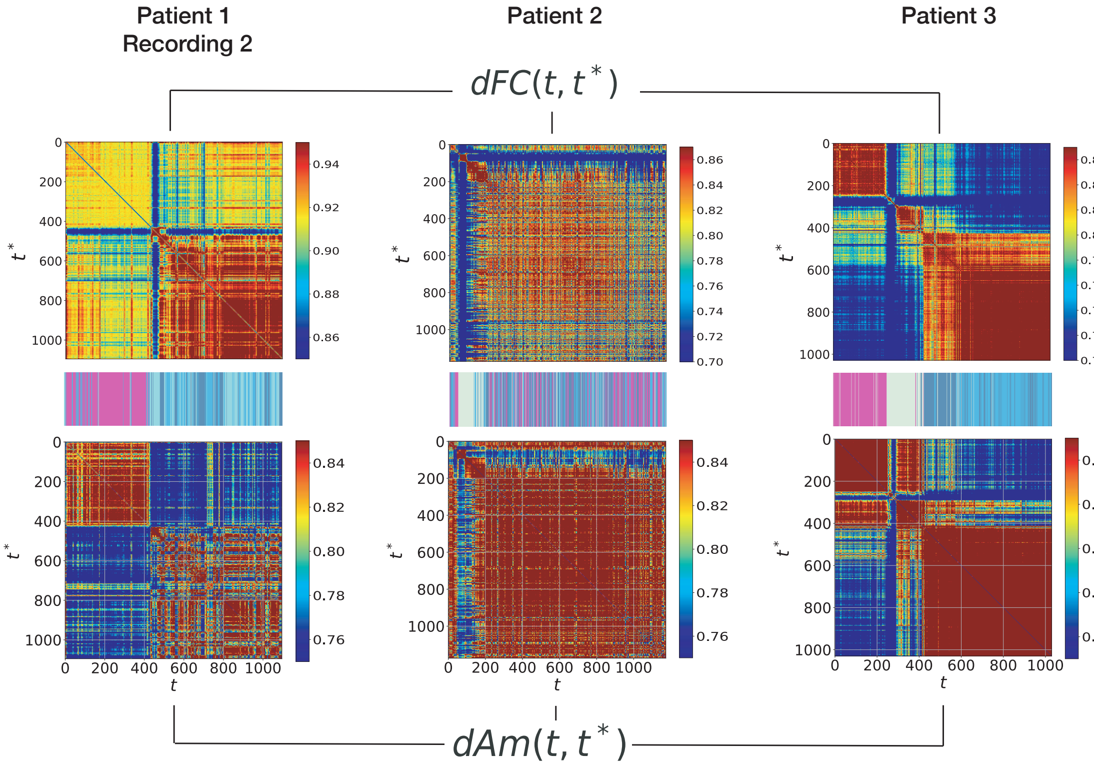


**Figure 6.** The **dFC** and **dAm**, as presented in **Figure 4.a** in the main text, computed for the 40Hz band of the other three recordings analysed. The barcode in between the two matrices, in each layer, represents the sequence of allegiance states. We note the presence of the four states - baseline magenta),seizure (aquamarine), recovery 1 (light blue) and recovery 2 (dark blue) - in all three recordings..

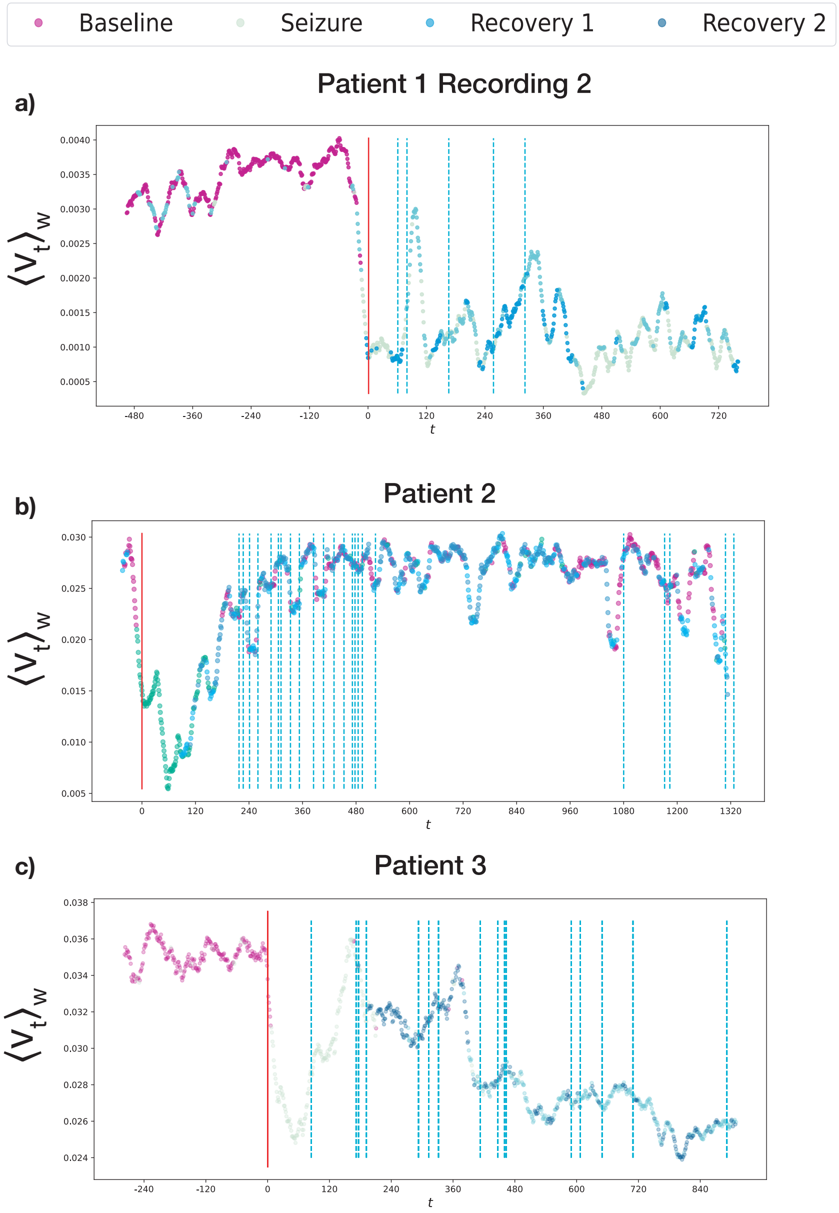


**Figure 7.** Each of the three plots corresponds to the analysis presented in **Figure 5.a** in the main text, for the other three recordings. In all three figures, each dot in corresponds to the average FC-speed value within a time window, colored with the color of the corresponding allegiance state - here it is evident that the baseline state (magenta points) is characterised by higher values of FC-speed, whereas the seizure state has the lowest values, hinting at a sudden slowing down of the overall network dynamic reconfiguration; recovery 1 points are concentrated in periods of rising FC-speed and on local maxima, whereas recovery 2 time windows happen around momentary slwoing downs of the dynamics and local minima.

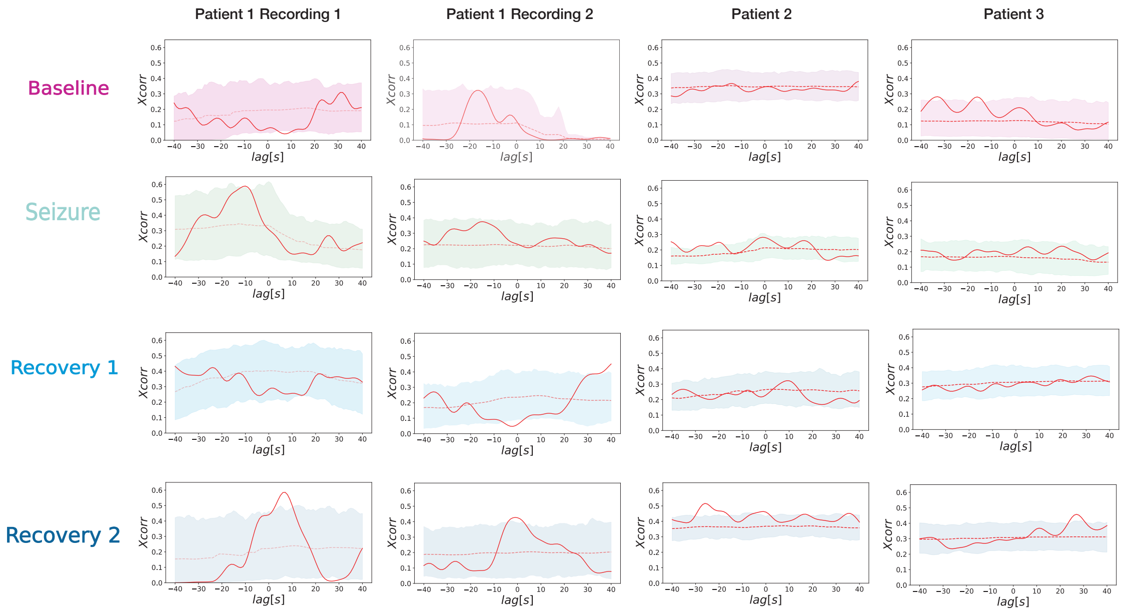


**Figure 8.** The red curves in the plots represent the lagged cross-correlation between time points of each allegiance state (baseline, seizure, recovery-1 and recovery-2) and the language dysfunction markers, as introduced in **Figure 5.c** in the main text; the shaded areas are comprised between the 5th and 95th percentiles of the values of cross-correlation computed for the time points of each state with a random permutation of the clinical markers in the [60s, 260s] interval.

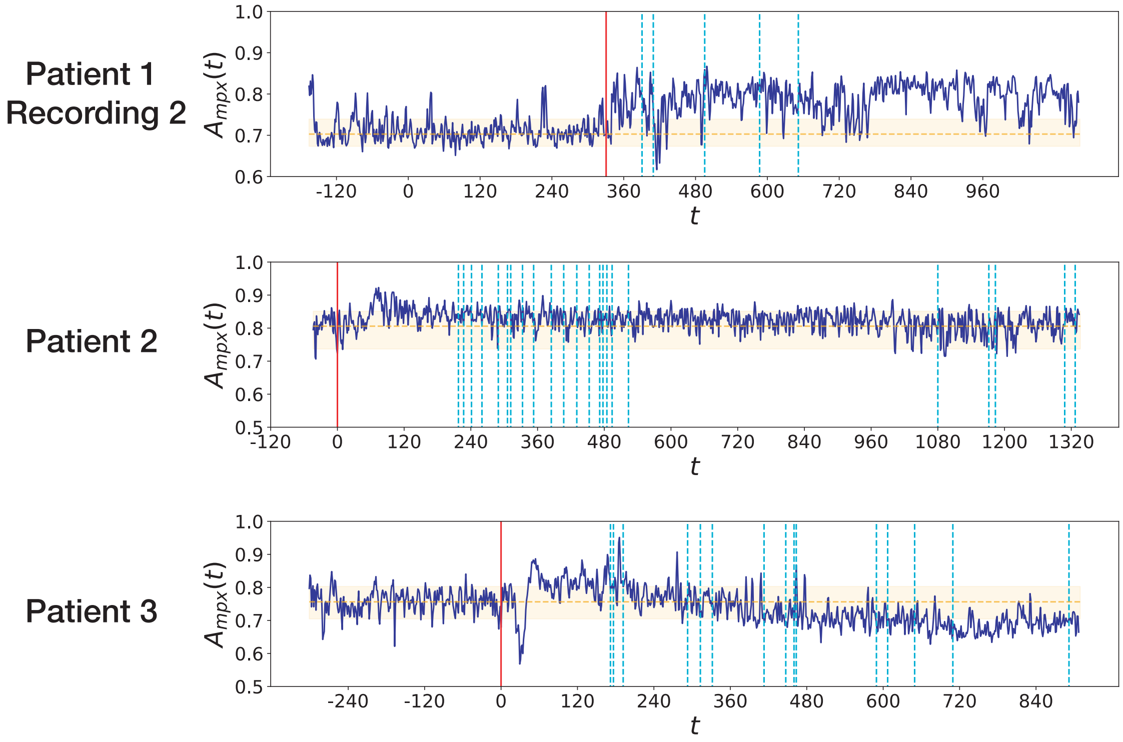


**Figure 9.** In each figure we plot the curve of inter-layer coallegiance (dark blue) and compare it to the average value of inter-layer coallegiance found in the baseline state (orange dotted line, the orange-shaded area corresponds to values between the 5th and 95th percentile of baseline inter-layer coallegiance values), as introduced in **Figure 6.b-d** in the main text for the first recording of Patient 1, computed for the other recordings.
